# Modulation of endogenous opioid signaling by inhibitors of puromycin sensitive aminopeptidase

**DOI:** 10.1101/2024.04.02.587756

**Authors:** Rohit Singh, Rongrong Jiang, Jessica Williams, Prakashkumar Dobariya, Filip Hanak, Jiashu Xie, Patrick E. Rothwell, Robert Vince, Swati S. More

**Affiliations:** Center for Drug Design, College of Pharmacy, University of Minnesota, MN, USA; Department of Neuroscience, University of Minnesota Medical School, Minneapolis, MN, USA

**Keywords:** . Puromycin sensitive aminopeptidase, puromycin aminonucleoside, enkephalin, analgesia, opioid receptor, metabolism, molecular modeling

## Abstract

The endogenous opioid system regulates pain through local release of neuropeptides and modulation of their action on opioid receptors. However, the effect of opioid peptides, the enkephalins, is short-lived due to their rapid hydrolysis by enkephalin-degrading enzymes. In turn, an innovative approach to the management of pain would be to increase the local concentration and prolong the stability of enkephalins by preventing their inactivation by neural enkephalinases such as puromycin sensitive aminopeptidase (PSA). Our previous structure-activity relationship studies offered the S-diphenylmethyl cysteinyl derivative of puromycin (**20**) as a nanomolar inhibitor of PSA. This chemical class, however, suffered from undesirable metabolism to nephrotoxic puromycin aminonucleoside (PAN). To prevent such toxicity, we designed and synthesized 5′-chloro substituted derivatives. The compounds retained the PSA inhibitory potency of the corresponding 5′-hydroxy analogs and had improved selectivity toward PSA. In vivo treatment with the lead compound **19** caused significantly reduced pain response in antinociception assays, alone and in combination with Met-enkephalin. The analgesic effect was reversed by the opioid antagonist naloxone, suggesting the involvement of opioid receptors. Further, PSA inhibition by compound **19** in brain slices caused local increase in endogenous enkephalin levels, corroborating our rationale. Pharmacokinetic assessment of compound **19** showed desirable plasma stability and identified the cysteinyl sulfur as the principal site of metabolic liability. We gained additional insight into inhibitor-PSA interactions by molecular modeling, which underscored the importance of bulky aromatic amino acid in puromycin scaffold. The results of this study strongly support our rationale for the development of PSA inhibitors for effective pain management.

## INTRODUCTION

The past decades have established the role of the endogenous opioid system in the regulation of pain, anxiety, depression, learning, memory, and gastrointestinal function. The system is ubiquitous throughout central and peripheral nervous systems and consists of oligomeric associations of receptor subtypes: the mu (µ), delta (δ), kappa (κ) opioid receptors and orphanin FQ/nociceptin receptor [1, 2]. Interactions of these receptor systems with their endogenous peptide ligands such as β-endorphin, dynorphin, enkephalin, and nociceptin with differing affinity and efficacy are responsible for the modulation of pain perception [3]. Exogenous opioids have been used successfully for clinical management of pain. The ability of such opioids to produce euphoria presents high risk of abuse and likelihood of developing opioid-use disorder (OUD) [4]. However, the need for pain relief is critical with an estimated over 51 million US adults suffering chronic pain symptoms, resulting in problematic prescription opioid rates [5]. Collectively, pain and opioid use pose tremendous societal costs, with pain-related health care and lost productivity exceeding $635 billion and opioid abuse-related health care, lost productivity, reduced quality of life, and life lost due to overdose exceeding $1.03 trillion annually [6]. Strategies to minimize the use of exogenous opioids or utilization of non-opioid pain alternatives thus require urgent attention. Attempts to disconnect undesirable effects of opioids from their potent pain-relieving properties have been mostly unsuccessful. One approach to reduce or eliminate opioid use altogether is by harnessing the body’s own ability to block pain, which is imparted by endogenous pain-relieving neuropeptides, but with regulation of reward-related behaviors [7, 8]. Among the opioid neuropeptides, enkephalins are known to modulate pain through MOR (µ-opioid receptor) and DOR (δ-opioid receptor), with slightly greater affinity for DOR [9]. Encoded by the proenkephalin (PENK) gene, located on exon 8 (gene ID: 5179), their role in multiple aspects of OUD such as sensitization, opioid reward and reinforcing properties is increasingly understood [10]. Cleavage of pro-enkephalin liberates peptides of varying length and sequences such as the pentapeptides Met- and Leu-enkephalins, the heptapeptide MERF (Met-enkephalin-Arg-Phe) and the octapeptide MERGL (Met-enkephalin-Arg-Gly-Leu) [11]. These opioid peptides share a common *N*-terminal sequence, Tyr-Gly-Gly-Phe-(Met/Leu), referred to as the “opioid motif.” The presence of Tyr and Phe is necessary for binding to the opioid receptors. Apart from their role in neuropsychiatric conditions, analgesia, and learning, they modulate respiration, blood pressure, embryonic development, angiogenesis, wound repair, and hepatoprotective mechanisms [7]. PENK knockout mice characterized previously by Konig et al. displayed hyperalgesia in response to painful stimuli, supporting the importance of these peptides in pain perception [12]. The effect of the opioid peptides is however short-lived due to rapid hydrolysis by two classes of enkephalin-degrading enzymes: aminopeptidases and carboxypeptidases. Puromycin-sensitive aminopeptidase (PSA) and aminopeptidase N (APN) cleave the Tyr-Phe bond, dipeptidyl aminopeptidase (DPP) hydrolyzes the Gly-Gly amide bond, while carboxypeptidases such as endopeptidase 24.11 (NEP) and angiotensin converting enzyme (ACE) act on the Gly-Phe bond (Figure 1) [13]. The amino terminal tyrosine is necessary for the opiate effects of these neuropeptides and thus degradation by aminopeptidases is considered the major pathway for the loss of their activity. Dual ENKephalinase Inhibitors (DENKIs) designed to inhibit APN and NEP have shown promise in preclinical testing and are being developed as alternatives to opiates in pain management [14, 15]. APN is localized exclusively to brain microvessels. The neural aminopeptidase PSA with the highest cytosolic and membrane expression is considered one of the major modulators of enkephalin activity [16]. The mice deficient in the PSA gene, generated by the gene trap method (Goku mice) exhibit reproductive defects as well as compromised pain response with increased anxiety [17]. The latter could result from changes in the levels of circulating enkephalins.

**Figure 1.**
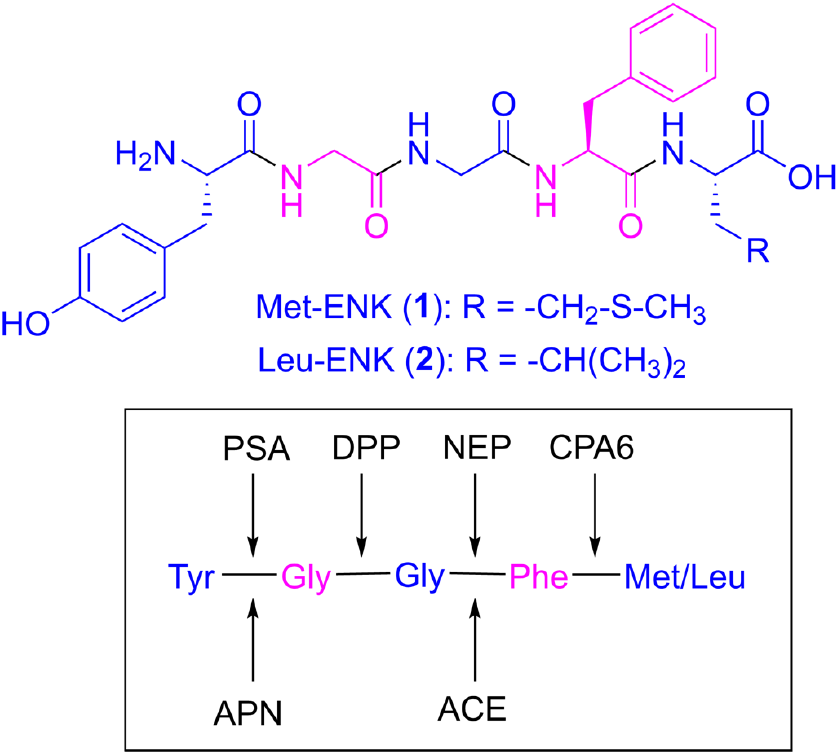
Chemical structures of Met-Enkephalin (**1**) and Leu-Enkephalin (**2**). Cleavage sites for proteolytic hydrolysis are depicted with arrows. **PSA**: Puromycin Sensitive Aminopeptidase; **APN**: Aminopeptidase N; **DPP:** Dipeptidyl Peptidase; **NEP**: Neprilysin; **ACE**: Angiotensin Converting Enzyme; **CPA-6**: Carboxypeptidase A-6.

PSA belongs to the M1 family of zinc metallopeptidases, which contains two histidines and a distal glutamate coordinating with a catalytic zinc(II) [18]. Apart from its role in enkephalin metabolism, it is also implicated in cell cycle control, proteosomal processing and development of anticancer agents. The expression of PSA in mouse brain has been correlated to tau [19, 20] and polyglutamine peptide levels [21] and resultant neurodegeneration. It plays a neuroprotective role in amyotrophic lateral sclerosis by providing a degradation pathway for neurotoxic proteins such as SOD1 [22]. Recently, a crystal structure of PSA was reported at 2.3 Å resolution, which explains the broad substrate tolerance of this enzyme [23]. The canonical inhibitor puromycin binds with the nucleoside portion interacting with the active site zinc and coordinating residues. Structural similarities between puromycin (**3**, Figure 2) and the aminoacyl adenyl terminal of aminoacyl tRNA allow puromycin to terminate protein synthesis, which has been exploited for the development of effective antibiotics [24]. However, the antibiotic efficacy of puromycin and related derivatives suffered due to their lack of selectivity toward pathogenic *vs*. host protein synthesis. We have previously explored the structure-activity relationship of puromycin-related compounds as antibiotic and anticancer agents with better selectivity and toxicity profiles [25, 26]. In recent years, we focused on enkephalinase inhibitory activity of puromycin for the development of analgesic agents. We reported upon the analgesic activity of an APN inhibitor, tosedostat, which was considered a clinical candidate for cancer [27]. Tosedostat synergized the analgesia of morphine, reducing effective opioid dose. Similarly, bestatin and thiorphan, inhibitors of aminopeptidase and endopeptidase respectively, potentiate the analgesic effect of Met-enkephalin after intracerebral administration [28, 29]. These studies, along with the success of DENKIs, support the use of enkephalinase inhibitors for achieving antinociceptive effect by modulating the levels of endogenous opioid peptides. Addressing the breakdown of endogenous neuropeptides could avoid overstimulation of opioid receptors and hence the development of dependence, addiction, or increased tolerance to analgesic effects.

**Figure 2.**
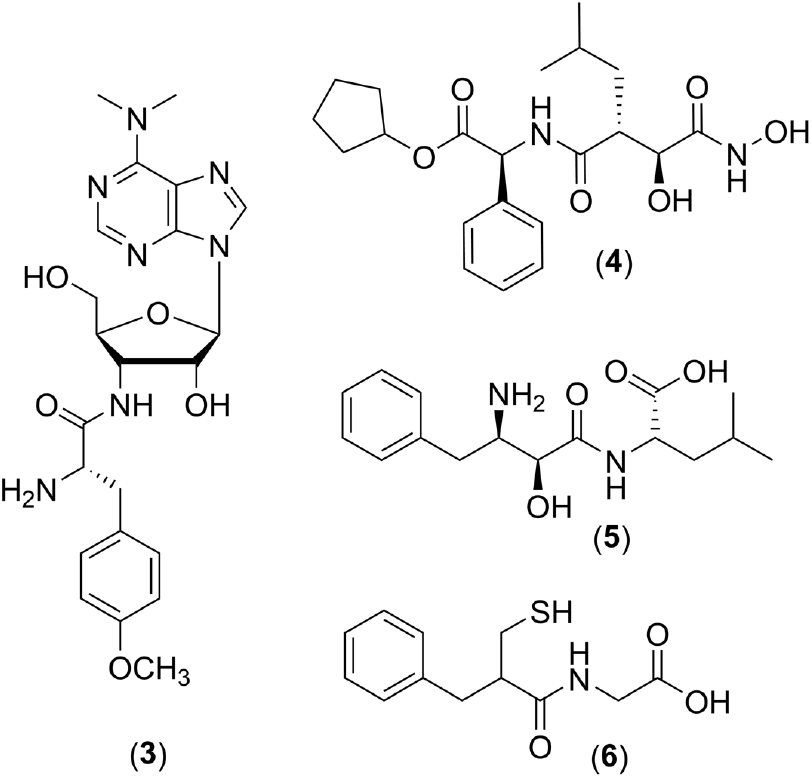
Known aminopeptidase inhibitors: Puromycin (**3**), Tosedostat (**4**), Bestatin (**5**) and Thiorphan (**6**).

Herein, we report the synthesis of lead puromycin analogs aimed at preventing the formation of nephrotoxic PAN. We also present exhaustive biochemical and in vitro evaluation of PSA inhibitors in support of our thesis. Antinociceptive effect of the lead compound was examined using the hot plate and tail flick assays in combination with Met-enkephalin. Finally, pharmacokinetic and computational studies were conducted to gain insight into metabolic liability and interactions with the target to improve PSA inhibitory potency. Collectively, the results of this study support the development of selective PSA inhibitors as a strategy for alternatives or conjugates to opioid pain medications and validates neural PSA as a therapeutic target.

## RESULTS AND DISCUSSION

### Chemistry

The puromycin aminonucleoside (PAN, **6**) resulting from the hydrolysis of puromycin is devoid of antibiotic activity, but is nephrotoxic in animals and humans [30]. PAN is monodemethylated by liver enzymes both in vitro and in vivo and subsequently converted to the 5′-nucleotide (**10**) [31]. The biosynthesis of the 5′-nucleotide (**10**) from monodemethyl-PAN (**9**) occurs in kidney slices. For this to happen, the demethylated PAN **9** from the liver enters the kidney, where nucleotide formation can occur. This nucleotide **10** is suggested to be the active metabolite of PAN which induces kidney toxicity. Thus, nephrotoxicity of puromycin has been ascribed to enzymatic hydrolysis of the methoxyphenylalanyl sidechain to release PAN (Scheme 1).

**Scheme 1.**
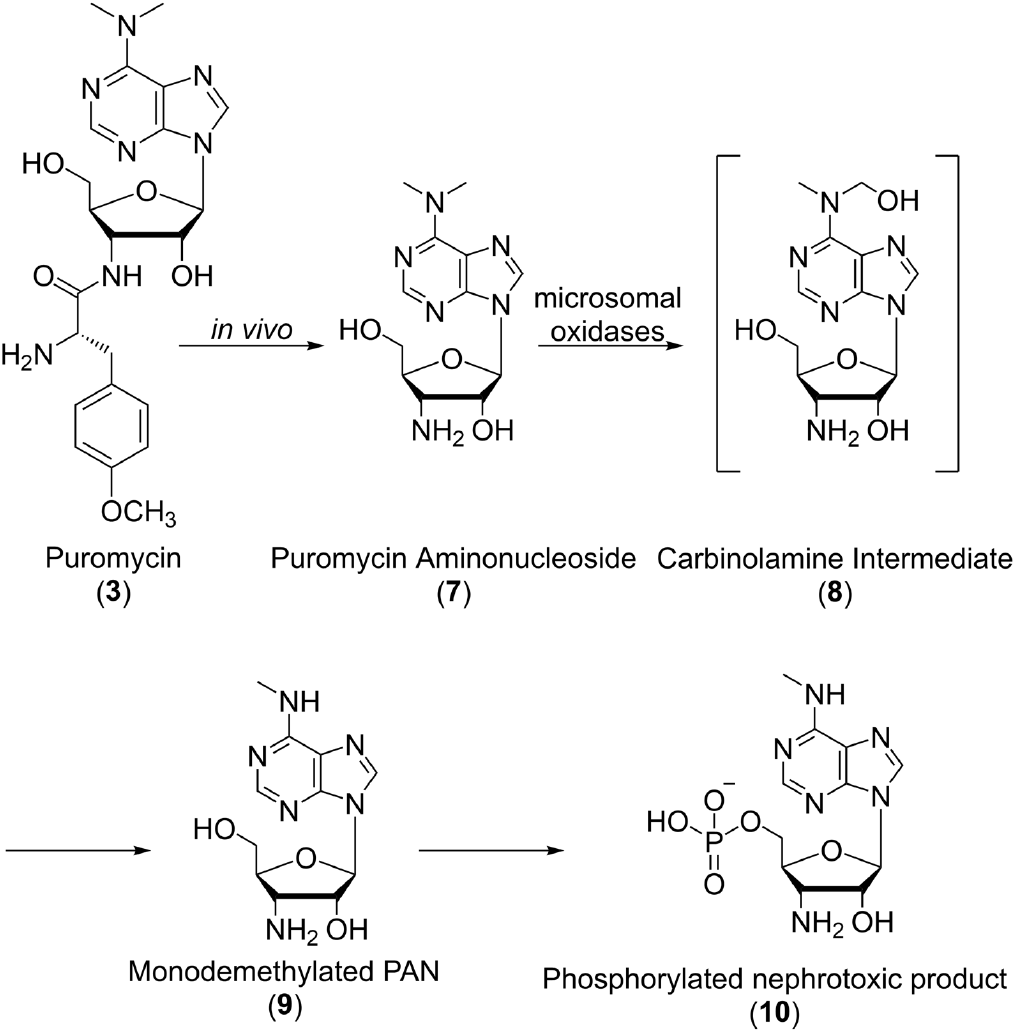
Mechanism of toxicity of puromycin

#### Synthesis of 5′-Substituted Inhibitors of Aminopeptidases

Our structure-activity relationship studies focused on eliminating the protein synthesis inhibitory property of the parent puromycin. We have previously demonstrated that non-aromatic aminoacyl substitution in puromycin structure obtained by replacing the aromatic 4-methoxyphenylalanine amino acid with cysteine could provide a handle to modulate such activity [26]. Specifically, S-alkyl substitutions with length up to six carbons retained protein synthesis inhibition, while larger aromatic substituents such as diphenyl and triphenyl were devoid of such activity. We thus incorporated a larger hydrophobic character in our design of new analogs. Synthesis of this series of compounds with 5′-hydroxy substituent was accomplished by coupling of PAN (**7**) with the appropriate *N*-Boc protected amino acids (**11**) (Scheme 2) [26]. Deprotection of *N*-Boc was achieved by anhydrous trifluoroacetic acid (TFA). Concerns about the potential nephrotoxicity of the liberated aminonucleoside metabolite still plagued this series of compounds. To design an active puromycin molecule that when hydrolyzed at the peptide bond would release a non-toxic aminonucleoside, we replaced the 5′-hydroxyl on the ribofuranosyl ring with chlorine. Such substitution of the sugar ring is known to confer resistance to conversion to the corresponding hydroxyl in both in vitro and in vivo studies, thus potentially removing the nephrotoxicity [32]. With PSA serving as the starting scaffold for the synthesis of these inhibitors, our previous structure-activity relationship studies guided the design of these inhibitors.

As outlined in Scheme 2, the substitution of the 5′-hydroxyl group in PAN (**7**) with a chloro group was achieved by halogenation in the presence of SOCl_2_ and triethyl phosphate to obtain compound **14** [32]. Coupling of the chlorinated intermediate **14** with N-Boc protected *S*-alkyl-L-cysteine derivatives (**11**) was conducted similarly to 5′-hydroxy derivatives using dicyclohexyl carbodiimide (DCC) and Nhydroxy succinimide (NHS), resulting in N-Boc protected analogs **15**. The N-Boc protecting group was removed within 10 minutes by anhydrous TFA providing the corresponding amine derivatives **16**. Possible hydrolysis of the glycosidic bond necessitates this transformation to be performed under strict anhydrous conditions. Purification of the final compounds was accomplished by in vacuo azeotropic removal of TFA with dry acetonitrile. The resultant residue was solubilized in methanol and passed through a column of freshly prepared (Cl^-^ to OH^-^) Amberlite resin. Finally, purification by flash chromatography, as detailed in Experimental Methods, furnished the desirable compounds (Figure 3). It is known that D-isomers of puromycin derivatives do not inhibit protein synthesis. To potentially exploit this property, a complimentary set of D-isomers about the *S*-diphenylmethyl residue was synthesized (Figure 3, compounds **24-25**).

**Scheme 2.**
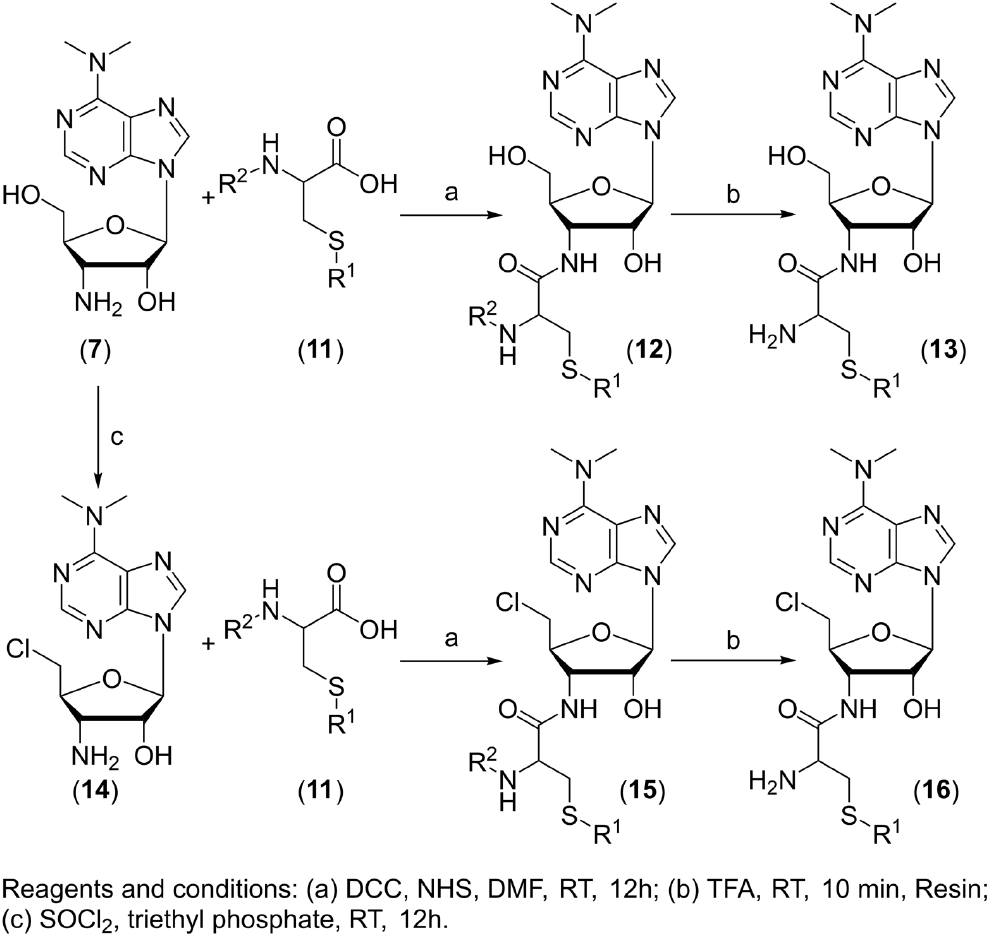
Synthesis of 5′-chloro substituted inhibitors of PSA

**Figure 3.**
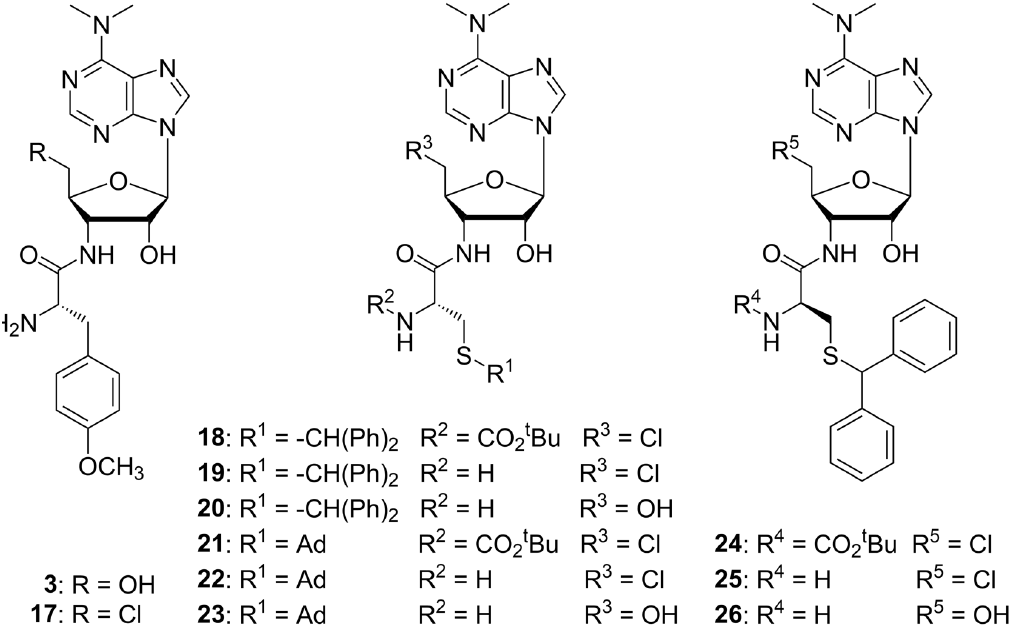
Structure activity study for the effect of variations on 5′ position of ribose ring

#### Dose-Response Studies Against Puromycin Sensitive Aminopeptidase (PSA) and Inhibitory Potency Against Aminopeptidase N (APN) and Neprilysin (NEP)

PSA is a neural metallopeptidase that hydrolyzes a single amino acid from the N-terminus of its substrate [33]. While the majority of the PSA protein (80%) is located within the cytoplasm, the remainder is associated with cell membranes [34]. Both cytoplasmic and membrane-associated forms are interchangeable and have been shown to degrade enkephalins in the brain [34]. Given the broad substrate specificity, PSA has been implicated in the regulation of cell cycle and pathological events in neurodegeneration and cancers [35]. We examined the inhibitory potency of the new puromycin analogs described here in PSA enzymatic assay (Table 1 and Figure 4). Selectivity toward PSA was determined by performing inhibitory assays with enzymes APN and NEP. Enzymatic activity of previous 5′-hydroxy series is included for a comprehensive structure-activity relationship (SAR) [26]. Enzymatic assays were performed using purified PSA protein expressed in our lab. The assay utilized alanine-4-methoxy-2-naphthylamide (Ala-4-MNA) as a substrate for the enzymatic reaction. Release of 4-MNA was monitored by excitation at 340 nm and emission at 425 nm for 30 min at room temperature.

**Figure 4.**
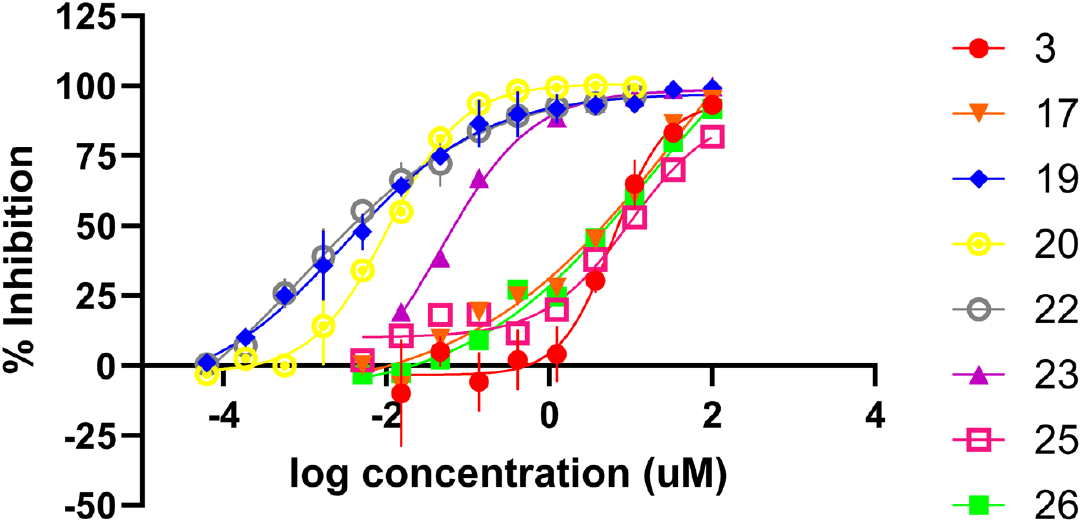
Dose–response curves for the inhibition of PSA

**Table 1.**
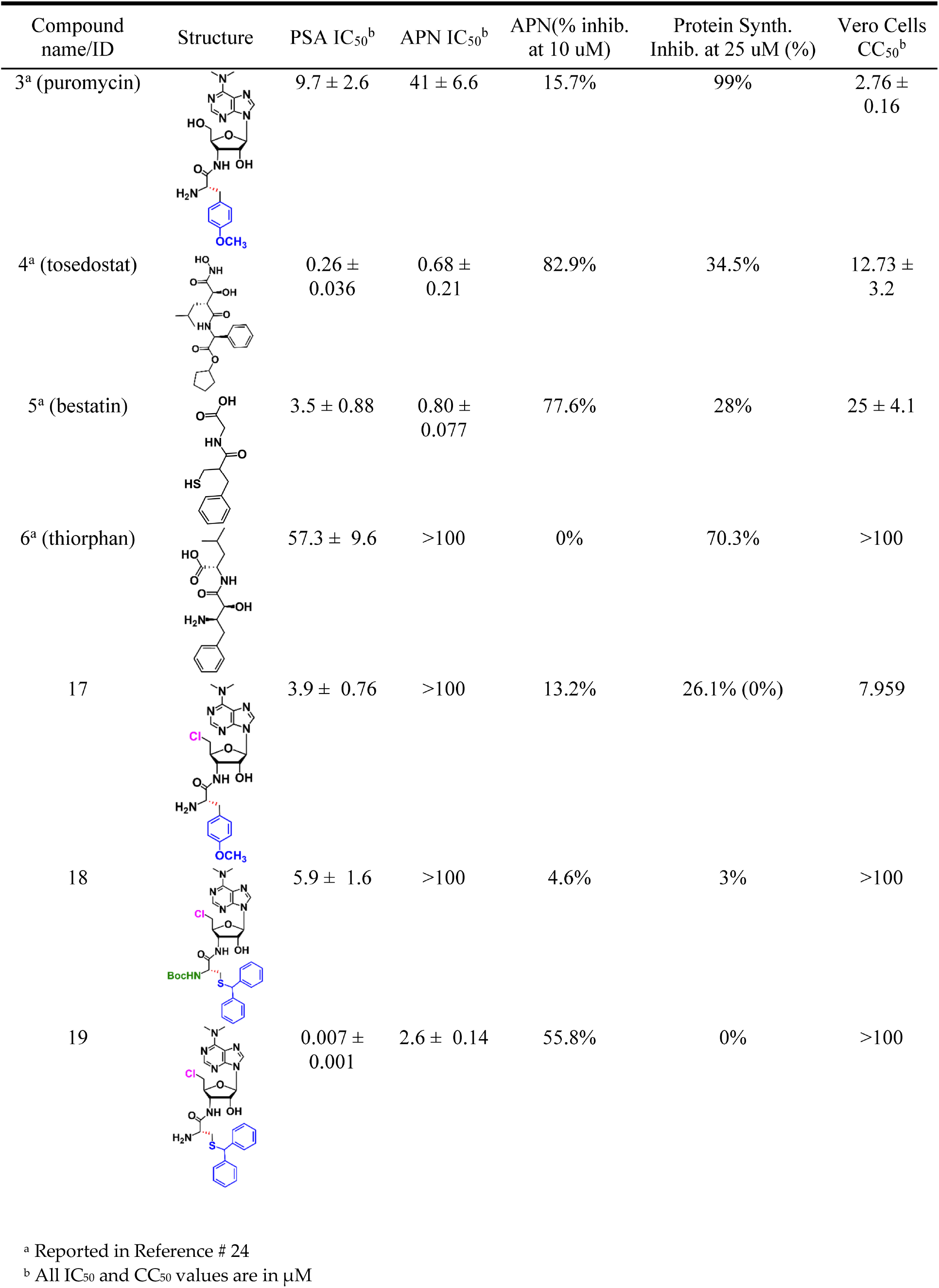
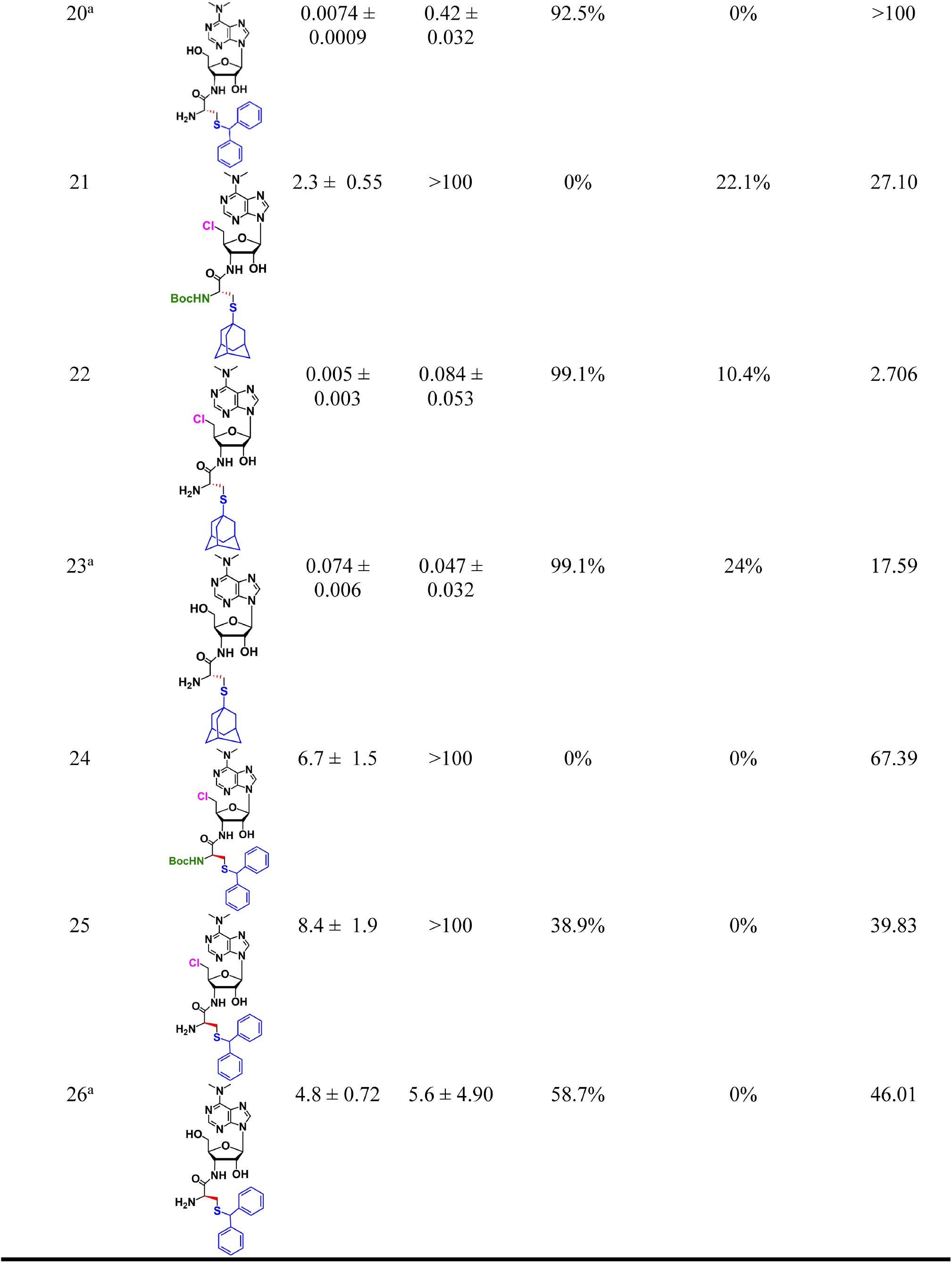
IC_50_ values for the inhibition of Puromycin Sensitive Aminopeptidase (PSA) and Aminopeptidase N (APN).

| Compound name/ID | Structure | PSA IC<sub>50</sub><sup>b</sup> | APN IC<sub>50</sub><sup>b</sup> | APN(% inhib. at 10 uM) | Protein Synth. Inhib. at 25 uM (%) | Vero Cells CC<sub>50</sub><sup>b</sup> |
| --- | --- | --- | --- | --- | --- | --- |
| 3<sup>a</sup> (puromycin) |  | 9.7 ± 2.6 | 41 ± 6.6 | 15.7% | 99% | 2.76 ± 0.16 |
| 4<sup>a</sup> (tosedostat) |  | 0.26 ± 0.036 | 0.68 ± 0.21 | 82.9% | 34.5% | 12.73 ± 3.2 |
| 5<sup>a</sup> (bestatin) |  | 3.5 ± 0.88 | 0.80 ± 0.077 | 77.6% | 28% | 25 ± 4.1 |
| 6<sup>a</sup> (thiorphan) |  | 57.3 ± 9.6 | >100 | 0% | 70.3% | >100 |
| 17 |  | 3.9 ± 0.76 | >100 | 13.2% | 26.1% (0%) | 7.959 |
| 18 |  | 5.9 ± 1.6 | >100 | 4.6% | 3% | >100 |
| 19 |  | 0.007 ± 0.001 | 2.6 ± 0.14 | 55.8% | 0% | >100 |

<sup>a</sup> Reported in Reference # 24
<sup>b</sup> All IC<sub>50</sub> and CC<sub>50</sub> values are in μM

|  |  |  |  |  |  |  |
| --- | --- | --- | --- | --- | --- | --- |
| 20<sup>a</sup> | | $0.0074 \pm 0.0009$ | $0.42 \pm 0.032$ | 92.5% | 0% | >100 |
| 21 | | $2.3 \pm 0.55$ | >100 | 0% | 22.1% | 27.10 |
| 22 | | $0.005 \pm 0.003$ | $0.084 \pm 0.053$ | 99.1% | 10.4% | 2.706 |
| 23<sup>a</sup> | | $0.074 \pm 0.006$ | $0.047 \pm 0.032$ | 99.1% | 24% | 17.59 |
| 24 | | $6.7 \pm 1.5$ | >100 | 0% | 0% | 67.39 |
| 25 | | $8.4 \pm 1.9$ | >100 | 38.9% | 0% | 39.83 |
| 26<sup>a</sup> | | $4.8 \pm 0.72$ | $5.6 \pm 4.90$ | 58.7% | 0% | 46.01 |

Among known inhibitors of aminopeptidases, puromycin (**3**) and bestatin (**5**) displayed low micromolar PSA inhibitory activity (IC_50_ 9.7 and 3.5 µM, respectively), while tosedostat was the most potent (IC_50_ 0.26 µM). Puromycin displayed 4.3-fold selectivity toward PSA than APN. Substitution of 5′-hydroxy with 5′-chloro improved selectivity, with a marginal increase in efficacy against PSA. Previous SAR informed about the importance of hydrophobic steric restrictions on amino acid residue in **3**, wherein diphenyl substitution on the cysteinyl sulfur resulted in more than a magnitude improvement in PSA inhibitory potency (IC_50_ 0.0074 ± 0.0009 µM for compound **20**). In this study, we examined the effect of 5′-chloro substitution on the inhibitory potency of the S-diphenylcysteinyl-puromycin derivative **20**. The 5′-chloro compound **19** (IC_50_ 0.0078 ± 0.0009 µM) retained the inhibitory potency of the parent 5′- hydroxy derivative **20** against PSA. Akin to puromycin, a 5′-chloro substitution improved selectivity toward PSA from 57-fold for compound **20** to 371-fold for compound **19**. Compound **19** was less active against APN compared to compound **20** (IC_50_ 2.644 for compound **19** *vs* 0.42 ± 0.032 µM for **20**), thus improving the therapeutic window. Both compounds were inactive against neprilysin (NEP). In accordance with our previous observation, the corresponding D-isomer of the 5′-chloro-S-diphenyl compound (compound **25**) was over one order of magnitude less active than the corresponding L-isomer in enzymatic assays with PSA, APN and NEP [36]. Interestingly, substitution of 5′-chloro in the D-isomer **25** also offered selectivity toward PSA over APN compared to the corresponding 5′-hydroxy derivative **26**. N-Boc protected analogues of compounds **19** and **25** were much less potent than the respective free amine derivatives. We then examined the effect of hydrophobic aliphatic substitution on cysteinyl sulfur by incorporating the bulky adamantyl group. The inhibitory potency of S-adamantyl substituted analog **22** was similar to S-diphenyl substituted derivative **19**. The compound, however, effectively inhibited APN with an IC_50_ 0.031 ± 0.0009 µM. Thus, the advantage in selectivity offered by 5′-chloro substituent in Sdiphenyl substituted cysteine derivatives was not seen in S-adamantyl compound **22**, suggesting the importance of aromatic substituent at this site for PSA activity. There was no inhibition of NEP observed in the presence of compound **22**. Based on this limited structure-activity relationship analysis, the 5′- chloro-S-diphenylcysteinyl derivative **19** appeared to be the most potent and PSA selective candidate and was employed in further in vitro and in vivo investigations.

#### In vitro Determination of Protein Synthesis Inhibitory Activity and Cellular Toxicity

Apart from preventing the formation of nephrotoxic PAN, our design rationale was to improve the therapeutic profile for this class of compounds. Thus, the design rationale based on puromycin structure needs to be thoroughly examined for its protein synthesis inhibitory activity, which is responsible for toxicity toward normal cells. However, differences in cellular update and binding of puromycin have also offered unrelated toxic effects [37, 38]. Toward that goal, we studied the ability of the synthesized 5′-chloro derivatives to inhibit protein synthesis as well as to produce cytotoxicity in kidney epithelial cells. The assay utilized the 1–Step Human In Vitro Protein Expression kit (Thermo Scientific) to study the inhibition of translation and post–transcriptional modification of the GFP control plasmid by PSA inhibitors [26]. The synthesized inhibitors were tested at 25 µM and puromycin was employed as a positive control. Notably, at the tested concentration, puromycin was responsible for 99% inhibition of protein synthesis. The most active compound **19** against PSA, however, did not affect protein synthesis, affording a lead with promising toxicity–profile. Bulky adamantyl substituted compound **22** showed modest inhibition of protein expression at 25 µM (10.4%).

The results of protein synthesis inhibition assay were validated by performing cytotoxicity evaluation of these compounds against Vero cells in an MTT assay. The most active compound against PSA, compound **19**, did not inhibit the viability of Vero cells and offered a CC50 value greater than 100 µM. This effect appears to stem from steric constraints of the bulky cysteinyl substituents for these compounds, which is responsible for dissociating the PSA inhibitory property from ribosomal activity. Amongst the known inhibitors of aminopeptidases, except for thiorphan, bestatin and tosedostat showed modest inhibition of Vero cell growth with CC_50_ in low micromolar range. The adamantyl derivative **22** was cytotoxic with CC_50_ of 2.706 µM. Surprisingly, the D-isomer of the S-diphenyl derivative, compound **25**, although not a protein synthesis inhibitor, was cytotoxic with a CC_50_ of 39.83 µM. While there could be several factors responsible for undesirable cytotoxicity in Vero cells, our experimental data excludes the possibility of the involvement of protein synthesis inhibition being the cause for this effect.

#### Pharmacological Studies for the Determination of Antinociceptive Effects

Attempts to enhance stability and thus pharmacological action of endogenous neuropeptides by inhibiting enkephalinases has gained attention in recent years. Efforts directed at inhibiting multiple such peptidases with an aim to produce significant analgesia have shown promise in preclinical studies [39]. Specifically, several studies simultaneously targeting APN and NEP which are involved in the breakdown of enkephalins have been reported. Such class of compounds is referred to as DENKIs (dual enkephalinase inhibitors) [40], are being advanced to clinical evaluation for their nociceptive effects [15]. Efforts toward the development of potent PSA inhibitors with analgesic activity are rather limited. The potent and selective PSA inhibitors identified through this investigation provide an opportunity to examine their antinociceptive property and consequently, establish the role of PSA in the degradation of opioid neuropeptides.

#### Tail flick assay

The tail-flick model represents a test of response to thermally induced high-intensity transient pain stimuli [41]. The tail-flick method mostly embodies a spinal reflex response to noxious stimuli. The time taken to either flick or withdraw its tail is recorded as a measure of the response. Figure 5A shows a time-course of analgesia after intracerebroventricular (i.c.v.) administration of compound **19**. When administered systemically by the intraperitoneal (i.p.) route, **19** did not cause substantial analgesia. This could potentially be due to metabolic instability or inefficient distribution in the central nervous system of this compound. Administration of compound directly into the brain by i.c.v. route caused significant analgesia, which was sustained over the first 45-minutes after the compound administration (Figure 5A). Attempted 100 and 50 µg doses of the compound offered a dose-dependent rise in the %maximum possible effect (MPE) over the vehicle control. The effect of i.c.v compound **19** on %MPE at first 15 minutes post administration is displayed in Figure 5B. A statistically significant rise in %MPE was noted over the corresponding vehicle group.

**Figure 5.**
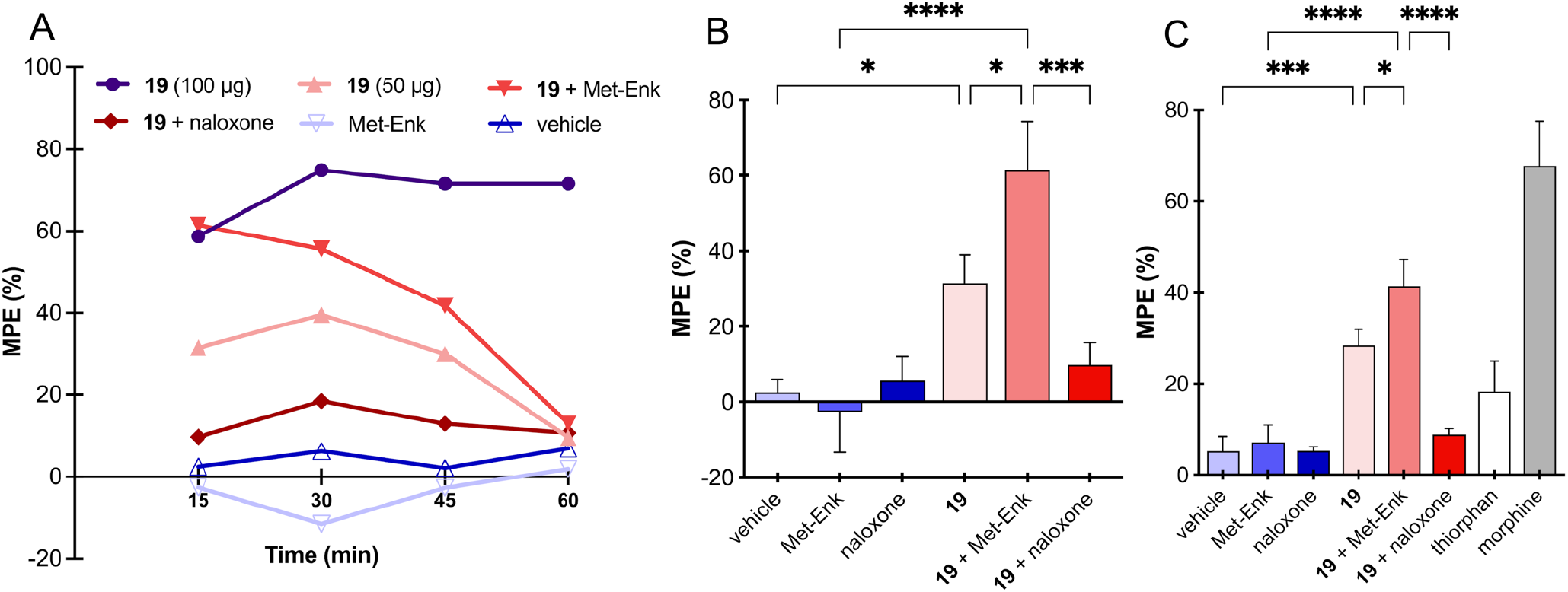
(A) Evaluation of time-dependent antinociceptive effects of compound **19** in tail-flick assay. The compound (50 or 100 µg/mouse) was administered intracerebroventricularly (i.c.v) alone or in combination with Met-Enk (50 µg/mouse). The combination with Met-Enk resulted in more pronounced anti-nociceptive effect compared to either compound alone. The increase in maximum possible effect (MPE) was inhibited by subcutaneous administration of opioid antagonist naloxone (5 mg/kg). The y-axis represents the percentage of maximum possible effect (%MPE). N=6-8/treatment group. Antinociceptive effect by tail-flick (B) and hot plate (C) assays at single time point (15 minutes) after compound administration are displayed (%MPE ± SEM). Statistical analysis was conducted by One-way ANOVA using Tukey’s multiple comparison test (* p < 0.05, *** p<0.001, **** p<0.0001).

**Figure 6.**
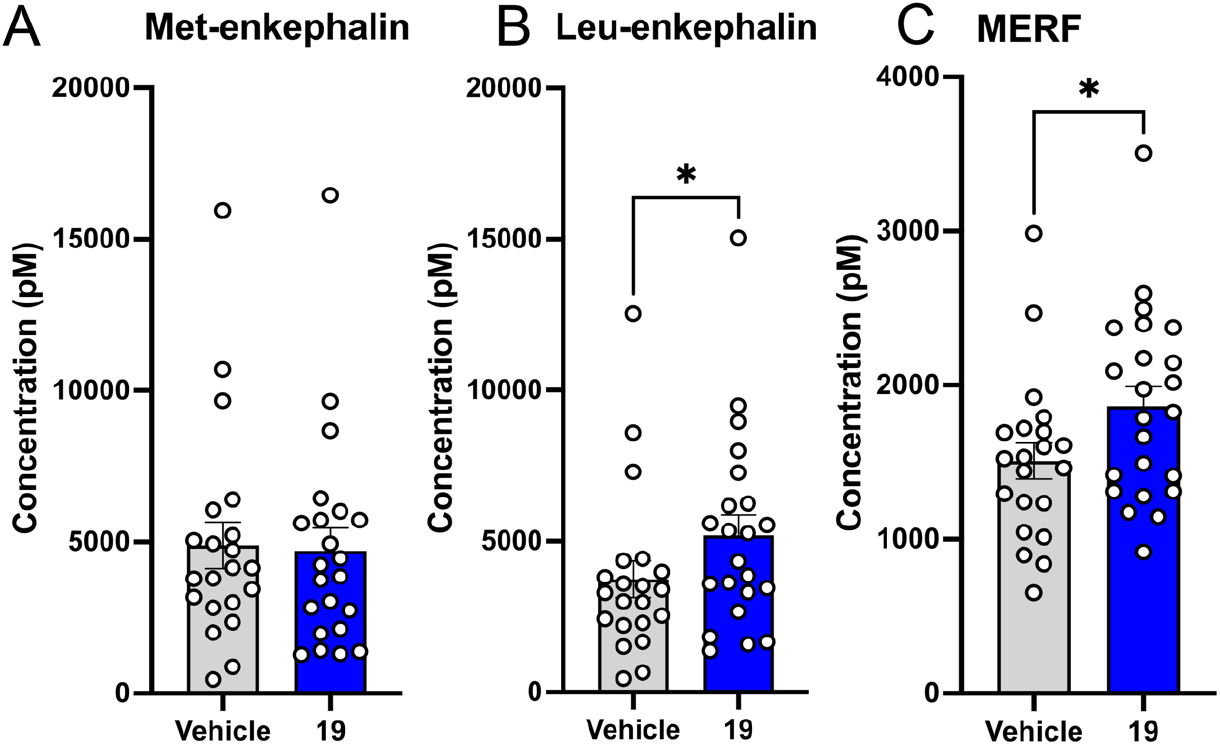
Effect of PSA inhibition by compound **19** on the release of endogenous enkephalins stimulated by KCl. Significant increase in the levels of Leu-Enk and MERF were detected by LC-MS/MS analysis as described in Methods. (N=6-8/treatment) Statistical analysis was conducted using nonparametric Mann-Whitney one-tailed U-test to determine the predicted increase in peptide levels (* p < 0.05).

#### Hot plate assay

In this transient pain model, a hot plate is utilized for the determination of antinociceptive potency of the lead compound against acute thermal stimulus [42]. This test is more sensitive to pain relay at the supraspinal level. Mice were administered compound **19** via i.c.v. route. Data collected at 15 minutes post compound administration is displayed in Figure 5C. Hot plate experiments corroborated our findings from the tail flick assay. Significant analgesic effect in animals injected intracerebroventricularly was observed in the experiment. In contrast to central administration, systemically delivered compound failed to register substantial analgesic response.

#### Effect of PSA inhibition on analgesia induced by exogenous Met-Enkephalin

This effort seeks to prevent metabolic degradation of endogenous enkephalins and thereby prolong their antinociceptive effect. By enhancing the body’s innate response to pain by increasing local concentrations of enkephalins where they are secreted, PSA inhibitors could foster analgesia with lesser side effects than morphine and other opiate drugs. Towards this end, we sought to determine how compound **19** would perform in the presence of Met-enkephalin (Figure 5). Co-administration of Met-enkephalin and compound **19** showed significantly improved %MPE above the compound-only groups in both tail flick and hot plate assays. To verify the involvement of opioid receptors in the observed antinociceptive activity reported here, we sought to study the effect of opioid receptor antagonist, naloxone, on these antinociceptive effect studies. Naloxone is a broad-spectrum opioid receptor antagonist known to interact with µ, δ, κ, and sigma (σ) opioid receptors, with the highest affinity for the µ receptors. The increase in the analgesic response of the combination was completely inhibited by administration of naloxone, a non-selective opioid receptor antagonist. This supports our hypothesis that the analgesic response mediated by PSA inhibitors and enkephalins is via their action on the opioid receptor. This result also indicates the potential of the combination of PSA inhibitors with opioid receptor agonists, thus achieving a substantial reduction in the latter’s effective dose while retaining the magnitude of the analgesic effect.

#### Effect of PSA inhibition on release and levels of enkephalins

The local release of enkephalin peptides is regulated by body’s innate pain response. However, the action of the peptides is controlled by proteolytic degradation by enkephalinases such as PSA. PSA has been implicated in the degradation of conventional Met-enkephalin or Leu-enkephalin in brain tissue. The Penk gene also encodes hepta- and octapeptides, MERF and MERGL [11]. These peptides have varying affinity and selectivity for opioid receptors and are processed by various enzymes. We have previously developed a liquid chromatography-tandem mass spectrometry (LC-MS/MS)-based assay for quantitation of extracellular levels of enkephalins and other neuropeptides released from mouse brain slices [43]. Treatment with KCl (50 mM) was used to stimulate the release of these peptides. In the presence of compound **19**, we observed significantly increased extracellular levels of Leu-enkephalin and MERF. Levels of Met-enkephalin, however, remained unchanged. This could potentially be due to the greater contribution of other enkephalinases such as APN, neprilysin and ACE, toward its degradation in the brain regions contained in the tissue slices. Our previous study demonstrated a similar effect of pharmacological inhibition of aminopeptidase N and neprilysin by bestatin and thiorphan combination on the release of Leu-Enk and other neuropeptides [43]. The results of this experiment support our rationale of enhancing analgesia by prolonging enkephalin stability and thus increasing resultant concentrations of these peptides by local PSA inhibition.

#### In vitro metabolic stability in plasma and liver microsomes

This study involved a strategy to prevent the formation of nephrotoxic PAN, resulting in compounds with no protein synthesis inhibitory activity and retention of the beneficial PSA inhibitory potential. To validate our rationale about improving the therapeutic profile of puromycin derivative by the inclusion of 5′-chloro substituent, we performed extensive stability assessment in in vitro model systems. Metabolite identification was conducted to verify the formation of 5′-hydroxyamino nucleoside (PAN).

#### Plasma Stability Studies

For in vitro plasma stability assessment, compound **19** was incubated in mouse and human plasma and aliquots collected at various time points were analyzed by LC-MS/MS to quantitate concentration of the parent compound with respect to time zero [44]. Compound **19** was stable in both mouse and human plasma during a 24 h incubation period at 37 °C (Table 2). Robust stability in plasma indicated resistance of the amide link in this compound to cleavage by serum peptidases.

**Table 2.** In vitro mouse and human plasma stability of 19.

| Time<br>(h) | % Remaining |  |  |  |
| --- | --- | --- | --- | --- |
|  | Mouse |  | Human |  |
|  | Average | % RSD | Average | % RSD |
| 0 | 100.0 | 0.0 | 100.0 | 0.0 |
| 1 | 103.7 | 9.4 | 97.6 | 2.9 |
| 3 | 106.5 | 4.7 | 100.4 | 4.3 |
| 6 | 111.9 | 11.0 | 102.3 | 8.3 |
| 24 | 116.7 | 4.0 | 109.0 | 3.4 |

#### Liver Microsomal Stability Studies

With no significant metabolism detected in plasma incubations, we sought to determine the metabolic stability of our lead compound in liver microsomes [45]. Such a parameter is considered an important property for early drug discovery projects and is expected to highlight metabolic liability inherent in chemical structures. A majority of xenobiotics are cleared predominantly by metabolism through cytochrome p450 present in liver microsomes and the information gained would be useful in improving in vivo pharmacokinetic profile, systemic bioavailability and potentially addressing any unforeseen toxicity issues [46]. Microsomal stability of the lead compound **19** was evaluated by monitoring the disappearance of test compound over 30 min [44]. Incubations were performed in the presence and absence of reduced nicotinamide adenine dinucleotide phosphate (NADPH), which is essential for the catalytic cycle of CYP (Figure 7). The concentration of the compound with respect to time determined by LC-MS/MS was used to calculate the remaining percentage at each time point with respect to time zero. Verapamil was used as a control as it is metabolized extensively by cytochrome P450. The metabolic rates of verapamil tested in mouse and human liver microsomes were consistent with the values in the literature. The metabolic profile of compound **19** was comparable to that of verapamil (Table 3). The metabolic half-lives of compound **19** and verapamil were 8.9 and 7.5 min, respectively, in mouse liver microsomes. Compound **19** was stable in the absence of NADPH in both mouse and human liver microsomes. The calculated intrinsic clearance (CL_int_) of compound **19** was high (156 uL/min/mg protein) and matched that of verapamil (185.6 uL/min/mg protein) under similar experimental conditions. This indicated extensive phase I metabolism of compound **19** by liver microsomes.

**Figure 7.**
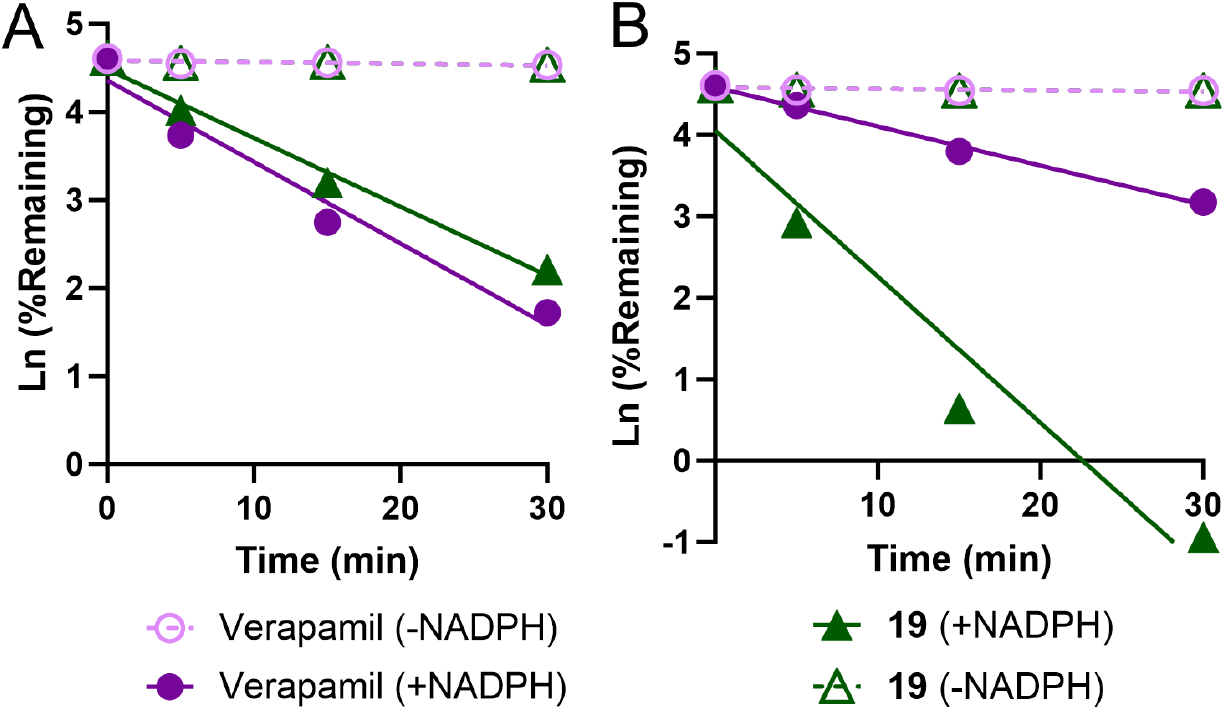
Time course of metabolic stability of **19** in mouse (A) and human (B) liver microsomes.

**Table 3.**
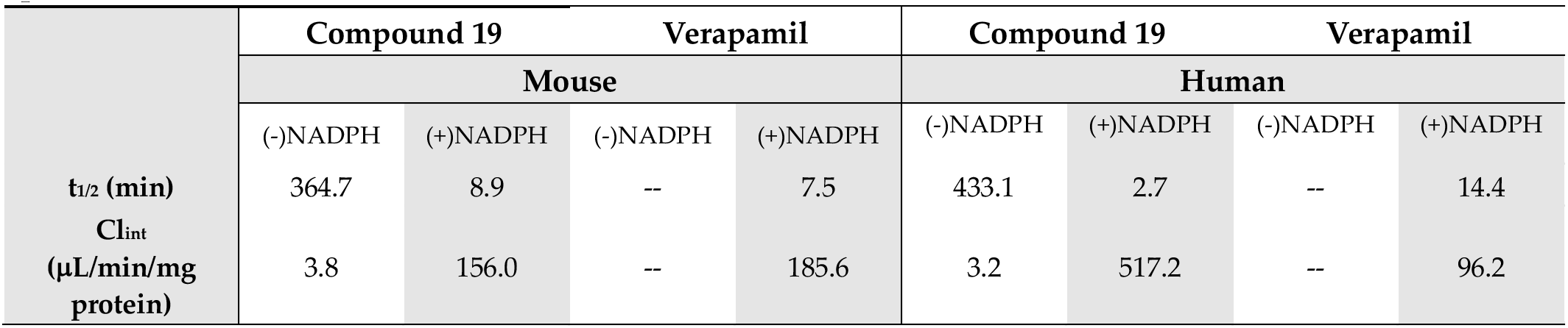
Calculation of in vitro pharmacokinetic parameters based on liver microsomal stability of compound 19.

Metabolite profiling was therefore performed in incubations with mouse and human liver microsomes. Figure 8 describes the postulated metabolites observed in the LC-MS/MS analysis. The oxidation of the sulfide in cysteine to the corresponding sulfoxide (**27**) contributed to 36% in human and 13.2% in mouse microsomes of all metabolites. Additionally, hydroxylation (mono-(**28**) and di-) of the aromatic ring(s) in the S-diphenylmethyl substituent, as well as N-oxidation of the cysteine amino group also made minor contributions. Oxidation of the primary amine (**29**) was also predicted. Human liver microsomal incubations of compound **19** identified reductive de-chlorination at the 5′-position to methyl group along with oxidation of the sulfide as a minor metabolite (**30**). N-demethylation of the purine ring (**31**) comprised 9.2% in mouse microsomes and is known to occur in the case of puromycin as well. None of the microsomal reactions yielded conversion of the 5′-chloride substituent to the corresponding 5′- hydroxy derivative, thus supporting our rationale. The results of this metabolic study would be valuable in designing the next generation of potent PSA inhibitors with better pharmacokinetic profiles.

**Figure 8.**
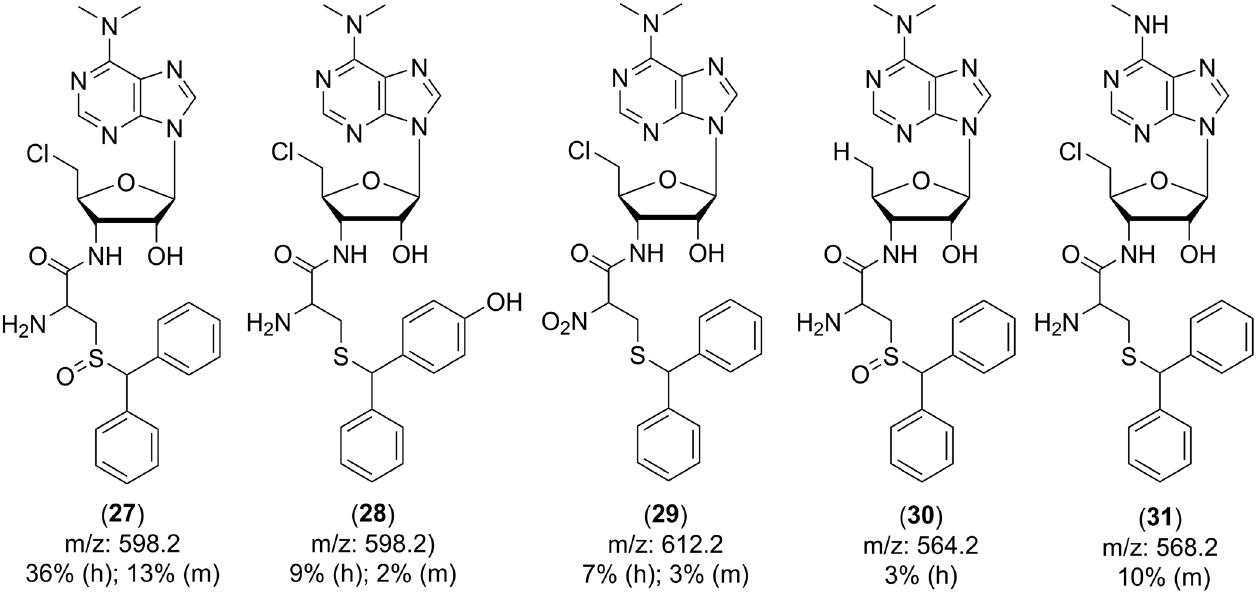
The postulated metabolites of **19** in mouse (m) and human (h) liver microsomes

#### Molecular Modeling and Docking Studies

To gain a better understanding of the PSA inhibitory results, molecular modeling analysis was performed by docking the lead compounds **19** and **22** into the active site of PSA. The X-ray crystal structure of human PSA (PDB code 8SW0) was recently reported by Madabushi et al. [23] and was imported for our ligand-receptor analysis. Validation of the receptor grid generated for docking studies was performed by docking the known ligand, puromycin (Figure 9A).

**Figure 9.**
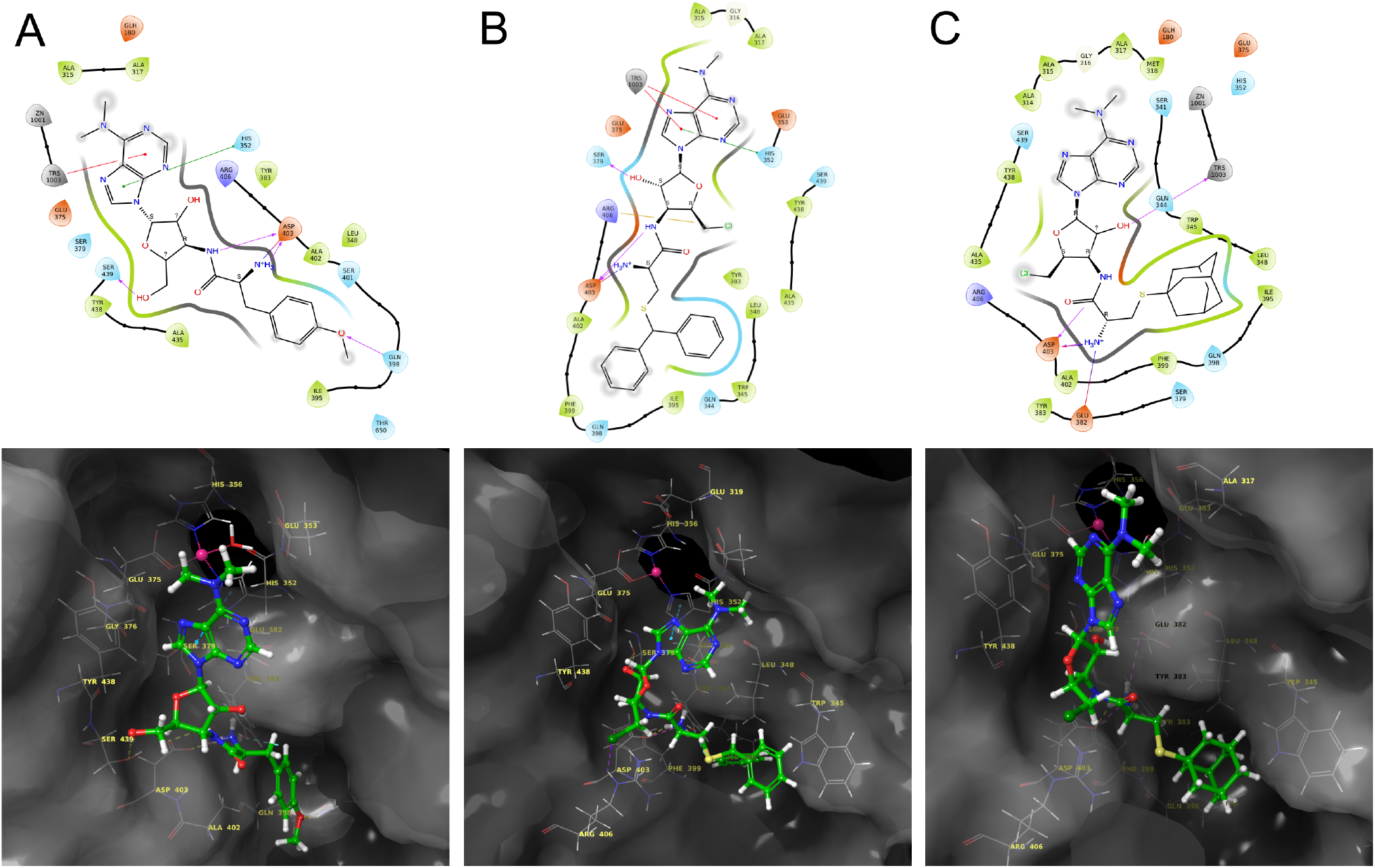
Molecular docking analysis of puromycin (A) and compounds **19** (B) and **22** (C). Top row shows 2D images of interactions of the inhibitor with PSA active site Zn(II) and surrounding residues. Bottom row shows fitting of each inhibitor within the binding site. Presence of Zn(II) and negatively charged residues direct the purine ring orientation, while accommodation of the aromatic amino acid residue in the hydrophobic pocket contributes favorably toward binding of this ligand alignment with the receptor.

The active site of PSA appears to be a long cleft with a catalytic site present at the closed end that contains the negatively charged residues Glu319 and Glu180, which correctly position the N-terminal end of the substrate. The long groove formed by hydrophobic and aromatic residues from helices α5 and α6 presents a rather large surface that likely is the reason for the broad substrate specificity of this enzyme [23]. All three low energy docked conformers of puromycin positioned the nucleoside portion in the proximity of the active site zinc(II) and the coordinating residues Glu375 at 1.96 Å, His352 at 2.12 Å, and His356 at 2.11 Å [23]. When docked, the dimethylamino (as ammonium) group of puromycin lies in proximity to the negatively charged pocket of the active site (Glu180 at 5.27 Å, Glu319 at 5.73 Å) and also interacts with the side chains of amino acids located in helix 11, helix 5, and helix 6. Notable π -π interaction occurs between the purine nucleobase and His352 (helix 5) and Tyr383 (helix 6). Hydrogen bonds (H-bond) are formed between 4’-amino of ribose in puromycin with Asp403 (helix 8) and H-bond of *p-* methoxy of the tyrosine residue with Gln398, all contributing toward the ligand-receptor complex. A salt bridge between the cationic α-ammonium group of the tyrosine residue in puromycin (**3**) and the anionic carboxylate of Asp403 also contributed significantly toward binding energy (docking score -8.390 kcal/mol). These results are consistent with the docking results of puromycin reported previously [23, 47].

Similar to puromycin, docking study revealed that the nucleoside portion of compound **19** and **22** positioned near the active site Zn and the coordinating residues (Figure 9B, 9C). Compound **19**, but not **22**, formed π -π stacking interactions with His352, resulting in favorable interaction with the metal center. Additionally, guanidinium group of the Arg406 formed ionic interactions with negative chloride substituent of compound **19**, which was absent in compound **22**. The S-diphenylmethyl substituent fits snuggly in the hydrophobic pocket formed by Phe399, Ile395, Trp 345, Leu348 and Tyr 383. This resulted in a favorable docking score for compound **19** (docking score -7.578 kcal/mol) over compound **22** (docking score -6.060 kcal/mol). A salt bridge between the ammonium of cysteine and carboxylate of Asp403 was present in compound **19**, similar to that formed by puromycin. In the case of compound **22**, this interaction was noted with Glu382.

Replacement of the 5′-chloride substituent by hydroxyl in compound **19** did not significantly affect the binding energy and the resultant docking score (-8.050 kcal/mol for compound **20**). Both compounds attained similar alignment and retained most of the active site interactions (Supporting Information, Figure S1). However, in the case of compound **22**, this substitution resulted in two distinct conformations of the purine ring; in proximity of Zn(II) or away from the metal center (Supporting Information, Figure S2). The alignment of nucleobase near Zn(II) offers a binding score similar to the 5′-chloro analogue **22** (docking score -5.26 kcal/mol), while conformation wherein purine prefers orientation toward hydrophobic pocket and formed a salt bridge between Glu375, resulted in improved docking score (-7.212 kcal/mol). The 5′-hydroxyl in compound **23** formed H-bond with Asp403. Additionally, salt bridge interaction of the parent chloride analog **22** was conserved (ammonium of compound **23** with Glu375 which is coordinated to Zn ion). Applying Zn as a constraint during docking resulted in slightly different alignments for puromycin and compound **19** (Supporting Information, Table S1) with changes noted in lig- and-receptor interactions. However, we did not observe any significant change in the binding energy and the resulting docking scores for these compounds. These *in silico* analyses support our experimental results for the observed PSA inhibitory activity of these compounds and therefore provide useful information for further design of potent PSA inhibitors.

## CONCLUSIONS

This study describes potent inhibitors of aminopeptidase PSAs based on the puromycin-based structural template identified in our previous study. The rationale for the design was based on the knowledge about the nephrotoxicity of the liberated PAN metabolite from structurally similar analogs. The 5′-chloro substituent on the sugar ring of the puromycin nucleoside was incorporated into the structure and the resulting compounds were studied for their inhibitory potency against aminopeptidases, PSA and APN. Compound **19**, the 5′-chloro derivative of puromycin coupled to S-diphenylmethyl-L-cysteine, retained PSA inhibitory activity and showed selectivity for PSA over other compounds tested. The steric bulk on the cysteine residue was responsible for better safety profile, with compound **19** being devoid of protein inhibition and cytotoxic potential. In vivo studies in mice demonstrated the analgesic potential of the PSA inhibitor and its ability to augment and prolong Met-enkephalin activity. The reversal of antinociceptive activity by naloxone confirms the involvement of opioid receptors in the pharmacological effect of the PSA inhibitor and Met-Enk combination. Compound **19** increased the endogenous release of Leu-Enk and MERF from brain slices, further supporting our rationale. Stability studies in plasma and liver microsomes did not indicate the conversion of the 5′-chloro substituent to the corresponding hydroxy derivative, which is important to prevent the formation of PAN. Collectively, the pharmacological results of this study along with *in silico* understanding of the binding of compound **19** to PSA, have offered insights into the design of the next generation of PSA inhibitors with better efficacy and safety profiles. This research effort provides a novel avenue for designing puromycin based aminopeptidase inhibitors, with the effective address of the dose-limiting toxicity of the parent compound. Early-stage stability assessment has revealed metabolic liabilities in this chemical class, which are currently being addressed in our newer generation of PSA inhibitors. Buttressing the body’s innate pain management system for the development of safer, non-addictive and effective is attractive in combating the opioid epidemic rampant in the US and worldwide. As we have shown in our previous study with tosedostat, the efficacy of aminopeptidase inhibitors can be augmented by supplementing with opioids, in effect reducing the effective doses of opioids and avoiding the emergence of tolerance and addiction. Further structural optimization to achieve desirable drug-like properties is thus clearly warranted to explore this promising avenue for pain management.

## EXPERIMENTAL SECTION

### Chemistry

#### General Procedures

All commercial chemicals were used as provided unless otherwise indicated. Whenever required, dry solvents were either purchased or dispensed from an anhydrous solvent system (J. C. Meyer) under argon. Teledyne ISCO’s Combiflash^®^ Rf system equipped with normal phase RediSep (silica) columns was used for flash chromatography. Moisture sensitive reactions were performed under an inert atmosphere of argon. Nuclear magnetic resonance (NMR) spectra were recorded on a Varian 600 MHz or a Varian 400 MHz spectrometer with Me_4_Si and residual chloroform or methanol as internal standards. Proton chemical shifts are reported in ppm as follows: integration, multiplicity, and coupling constant. The symbols for multiplicity are described as s = singlet, d = doublet, t = triplet, q = quartet, m = multiplet, br = broad, dd = double doublet, ddd = doublet of doublet of doublets, td = triplet of doublets, qd = quartet of doublets. High resolution mass spectra were recorded on Agilent TOF II TOF/MS instrument equipped with either an ESI or APCI interface. The analytical purity of samples was assessed on an Agilent 1260 HPLC with a UV detector using an Agilent Microsorb MV 100-5 C18 column (4.6 mm x 250 mm, 5 µM particle size). The chromatography conditions were as previously reported using a mobile phase of H_2_O (solvent A) and acetonitrile (solvent B) at 254 nm detector wavelength and flow rate of 1.0 mL/min. All tested compounds were >95% pure by HPLC analysis.

#### General procedure (A) for the coupling of amino acids with puromycin aminonucleoside

To a dry Schlenk tube purged with argon and equipped with a magnetic stir bar was loaded puromycin aminonucleoside (PAN, 50.0 mg, 0.17 mmol, 1 equiv.) and N-hydroxy succinimide (67.5 mg, 0.19 mmol, 1.1 equiv.). Appropriate N-protected amino acid substrate (0.19 mmol, 1.1 equiv.) was added to the flask under a cone of argon and the flask sealed with a rubber septum. Anhydrous dimethylformamide (DMF, 1.7 mL, 10 mL per mmol) was added to the reaction vessel via a syringe and the mixture stirred while being cooled to 0 °C in an icebath. Dicyclohexyl carbodiimide (DCC, 38.2 mg, 0.19 mmol, 1.1 equiv.) was then added to the reaction mixture and contents were stirred at 0 °C for 15 min. The ice bath was then removed, and reaction contents allowed to stir at room temperature for 18 h. The reaction progress was monitored by TLC analysis. After the completion of reaction, the precipitated dicyclohexyl urea side-product was filtered out using a Büchner funnel. The filter paper and the insoluble precipitates were washed with copious amounts of ethyl acetate. Filtrate was collected and removed in vacuo to yield the desirable product in crude form. Purification of the crude product was performed by flash chromatography utilizing Teledyne ISCO’s Combiflash^®^ Rf system equipped with normal phase RediSep (silica) columns, and mobile phase eluents as described below.

#### General procedure (B) for Boc-deprotection of amine

To a dry Schlenk tube purged with argon and equipped with a magnetic stirbar was loaded the N-protected starting material (1 equiv.) and the tube sealed with a rubber septum. An excess of anhydrous trifluoroacetic acid (TFA, 12 mL/mmol) was added to the reaction vessel via a syringe and the contents stirred at room temperature for 10 min under argon. After 10 minutes, the volatiles were removed in vacuo. To ensure complete removal of TFA, the contents in the reaction tube were azeotroped thrice with dry acetonitrile (1 x volume of TFA used) to obtain the product in crude form. A resin column of Amberlite IRA-410 OH^‒^ resin was freshly prepared as described:

Amberlite IRA-410 (20-25 mesh) Cl^‒^ form was (5 gm per 100 mL of sample solution) loaded on a column. 1 N solution of NaOH was prepared fresh by dissolving the appropriate amount of low chloride NaOH in deionized water and passed through the column (20 x volume of resin). The exchange of Cl^‒^ with OH^‒^ ions was monitored by the HNO_3_/AgNO_3_ test for the presence of Cl^‒^ ions in the eluent. Once the Cl^‒^ ions were undetectable in the eluent, deionized water was passed through the column and the pH of the eluent monitored by pH paper until it came down to 9. Next, the column was equilibrated by passing methanol through it (3 x volume of resin). The residue from the reaction vessel was redissolved in methanol and passed through this freshly prepared Amberlite IRA-410 OH^‒^ resin column. The column was washed with copious amount of methanol (10-15 x volume of resin). Removal of methanol in vacuo furnished the product in crude form. Purification of the crude product was performed by flash chromatography utilizing Teledyne ISCO’s Combiflash^®^ R_f_ system equipped with normal phase RediSep (silica) columns, and mobile phase eluents as described below.

#### General procedure for 5′-chlorination of PAN

To a Schlenk flask purged with argon was added PAN (206.4 mg, 0.7 mmol, 1 equiv.). Addition of triethyl phosphate (0.7 mL, 10 mL/mmol, excess) resulted in formation of a suspension. Thionyl chloride (203 µL, 2.8 mmol, 4 equiv.) was added and contents were stirred at RT for 16 hours under argon. The reaction was then poured into 100 mL of anhydrous Et2O, and allowed to stand for 1 h at RT. The resulting solid was collected by filtration and dissolved in methanol and passed through Dowex 1 resin (25-50 mesh) column freshly changed to –OH form by 1N NaOH. The ion-exchange column was washed with methanol (3 column lengths). Removal of MeOH in vacuo provided white powder, which was purified by flash chromatography utilizing Teledyne ISCO’s Combiflash® Rf system equipped with normal phase RediSep (silica) columns, and CHCl3/MeOH as mobile phase eluents to obtain compound **14** as white powder (148.4 mg, 68% yield).

Tert-butyl ((R)-3-(benzhydrylthio)-1-(((2S,3S,4R,5R)-2-(chloromethyl)-5-(6-(dimethylamino)-9H-purin-9-yl)-4-hydroxytetrahydrofuran-3-yl)amino)-1-oxopropan-2-yl)carbamate **(18)** The crude product obtained by following general procedure A as described above, was purified by flash chromatography utilizing Teledyne ISCO’s Combiflash^®^ R_f_ system equipped with normal phase RediSep (silica) columns with methanol/chloroform eluent mixture (neat CHCl_3_ for 10 min, to 20% MeOH/CHCl_3_ gradual gradient over the next 29 min) furnished compound **18** as white powder in 156.8 mg (79%) yield; R*_f_* 0.42 (MeOH:CHCl_3_/1:19); **^1^H-NMR** (600 MHz, CD_3_OD): 8.210 (2H, d, J = 4.00 Hz), 7.438 (4H, d, J = 2.29 Hz), 7.300 (4H, s), 7.23-7.21 (2H, m), 6.056 (1H, s), 5.34-5.33 (1H, m), 4.76-4.73 (1H, m), 4.599 (1H, t, J = 2.710 Hz), 4.282 (2H, t, J = 0.75 Hz), 3.912 (1H, dd, J = 12.34, 0.30 Hz), 3.776 (1H, dd, J = 11.62, 4.75 Hz), 3.490 (6H, s), 2.78-2.73 (1H, m), 2.64-2.60 (1H, m), 1.433 (9H, d, J = 2.87 Hz); **^13^C-NMR** (101 MHz, CD_3_OD) 172.28, 154.74, 152.86, 149.54, 141.21, 137.16, 128.22, 128.15, 128.05, 126.93, 119.93, 90.33, 81.57, 79.60, 73.77, 53.94, 53.59, 52.18, 44.21, 37.64, 33.66, 27.27; ESI-MS *m/z* [M + H]^+^ 682.2. (R)-2-amino-3-(benzhydrylthio)-N-((2S,3S,4R,5R)-2-(chloromethyl)-5-(6-(dimethylamino)-9H-purin- 9-yl)-4-hydroxytetrahydrofuran-3-yl)propenamide **(19)** The crude product obtained by following general procedure B as described above, was purified by flash chromatography utilizing Teledyne ISCO’s Combiflash^®^ R_f_ system equipped with normal phase RediSep (silica) columns with methanol/chloroform eluent mixture (neat CHCl_3_ for 8 min, to 20% MeOH/CHCl_3_ gradual gradient over the next 31 min) furnished compound **19** as white powder in 54.7 mg (41%) yield; R*f* 0.35 (MeOH:CHCl_3_/1:19); **^1^H-NMR** (600 MHz, CD_3_OD): 8.230 (2H, s), 7.440 (4H, dd, J = 10.74, 7.53 Hz), 7.298 (4H, q, J = 7.95 Hz), 7.209 (2H, q, J = 7.83 Hz), 6.075 (1H, d, J = 1.89 Hz), 5.302 (1H, s), 4.765 (1H, dd, J = 8.68, 5.82 Hz), 4.609 (1H, dd, J = 5.83, 1.85 Hz), 4.34-4.29 (1H, m), 3.936 (1H, dd, J = 5.83, 1.85 Hz), 3.803 (1H, dd, J = 12.29, 5.20 Hz), 3.60-3.40 (6H, m), 2.725 (1H, dd, J = 13.23, 6.89 Hz), 2.623 (1H, dd, J = 13.20, 6.47 Hz); **^13^C-NMR** (101 MHz, CD_3_OD) 174.80, 154.75, 151.88, 149.54, 141.49, 141.42, 137.16, 128.44, 128.15, 128.06, 128.03, 128.01, 126.86, 119.96, 90.37, 81.91, 73.79, 54.06, 54.03, 51.78, 44.32, 36.90, 31.32, 22.27; ESI-MS *m/z* [M + H]^+^ 582.2.

Tert-butyl ((R)-3-(((3S,5S,7S)-adamantan-1-yl)thio)-1-(((2S,3S,4R,5R)-2-(chloromethyl)-5-(6-(dimethylamino)-9H-purin-9-yl)-4-hydroxytetrahydrofuran-3-yl)amino)-1-oxopropan-2-yl)carbamate **(21)**

The crude product obtained by following general procedure A as described above, was purified by flash chromatography utilizing Teledyne ISCO’s Combiflash^®^ R_f_ system equipped with normal phase RediSep (silica) columns with methanol/chloroform eluent mixture (neat CHCl_3_ for 5 min, to 20% MeOH/CHCl_3_ gradual gradient over the next 19 min) furnished compound **21** as light-yellow powder in 123.3 mg (60%) yield; R*_f_* 0.48 (MeOH:CHCl_3_/1:19); **^1^H-NMR** (600 MHz, CD_3_OD): 8.226 (2H, d, J = 1.24 Hz), 6.074 (1H, s), 4.78-4.76 (1H, m), 4.623 (1H, d, J = 5.61 Hz), 4.34-4.33 (1H, m), 4.23-4.21 (1H, m), 4.029 (1H, d, J = 12.22 Hz), 3.879-3.86 (1H, m), 3.50-3.49 (6H, m), 2.93-2.90 (1H, m), 2.803 (1H, dd, J = 6.90, 6.11 Hz), 2.026 (3H, s), 1.880 (6H, s), 1.75-1.70 (6H, m), 1.448 (9H, s); **^13^C-NMR** (101 MHz, CD_3_OD) 172.29, 158.46, 154.74, 151.87, 149.55, 137.20, 125.04, 90.38, 81.63, 73.81, 54.95, 52.24, 44.29, 43.19, 35.89, 29.78, 27.27, 24.76; ESI-MS *m/z* [M + H]^+^ 650.2.

(S)-2-amino-3-(tert-butylthio)-N-((2S,3S,4R,5R)-5-(6-(dimethylamino)-9H-purin-9-yl)-4-hydroxy-2- (hydroxymethyl)tetrahydrofuran-3-yl)propenamide (**22)**

The crude product obtained by following general procedure B as described above, was purified by flash chromatography utilizing Teledyne ISCO’s Combiflash^®^ R_f_ system equipped with normal phase RediSep (silica) columns with methanol/chloroform eluent mixture (neat CHCl_3_ for 8 min, to 20% MeOH/CHCl_3_ gradual gradient over the next 30 min) furnished compound **22** as white powder in 36.2 mg (43%) yield; R*_f_* 0.40 (MeOH:CHCl_3_/1:19); **^1^H-NMR** (600 MHz, CD_3_OD): 8.153 (2H, d, J = 4.16 Hz), 6.003 (1H, s), 4.679 (1H, d, J = 2.48 Hz), 4.55-4.54 (1H, m), 4.265 (1H, s), 3.948 (1H, dd, J = 12.14, 2.06 Hz), 3.804 (1H, d, J = 5.06 Hz), 3.42-3.41 (6H, m), 2.799 (1H, dd, J = 12.10, 6.62 Hz), 2.651 (1H, dd, J = 12.43, 6.51 Hz), 1.944 (3H, s), 1.802 (6H, s), 1.642 (6H, s); **^13^C-NMR** (101 MHz, CD_3_OD) 174.27, 154.78, 151.91, 149.59, 137.20, 119.98, 90.43, 81.67, 73.85, 54.63, 54.61, 52.14, 44.19, 43.27, 39.42, 35.91, 35.89, 33.36, 29.82, 25.35; ESI-MS *m/z* [M + H]^+^ 550.2.

Tert-butyl ((S)-3-(benzhydrylthio)-1-(((2S,3S,4R,5R)-2-(chloromethyl)-5-(6-(dimethylamino)-9H-pu- rin-9-yl)-4-hydroxytetrahydrofuran-3-yl)amino)-1-oxopropan-2-yl)carbamate (**24**)

The crude product obtained by following general procedure A as described above, was purified by flash chromatography utilizing Teledyne ISCO’s Combiflash^®^ R_f_ system equipped with normal phase RediSep (silica) columns with methanol/chloroform eluent mixture (neat CHCl_3_ for 10 min, to 20% MeOH/CHCl_3_ gradual gradient over the next 29 min) furnished compound **24** as white powder in 39.8 mg (27%) yield; R*_f_* 0.42 (MeOH:CHCl_3_/1:19); **^1^H-NMR** (600 MHz, CD_3_OD): 8.123 (2H, dd, J = 8.70, 4.48 Hz), 7.339 (3H, dd, J = 15.80, 7.76 Hz), 7.190 (4H, dt, J = 17.74, 7.57), 7.14-7.08 (2H, m), 7.045 (1H, dd, J = 16.21, 8.53 Hz), 5.960 (1H, s), 5.226 (1H, s), 4.626 (1H, dd, J = 7.71, 6.58 Hz), 4.489 (1H, d, J = 5.05 Hz), 4.23-4.17 (2H, m), 3.850 (1H, dd, J = 12.31, 1.61 Hz), 3.77-3.71 (2H, m), 3.48-3.36 (6H, m), 2.72-2.69 (1H, m), 2.57-2.53 (1H, m), 1.365 (9H, s); **^13^C-NMR** (101 MHz, CD_3_OD) 172.32, 153.97, 151.88, 148.97, 141.34, 137.23, 128.17, 127.97, 126.87, 119.86, 90.02, 81.82, 73.85, 69.34, 53.90, 52.21, 44.23, 36.77, 33.67, 27.17; ESI-MS *m/z* [M + H]^+^ 682.2.

(S)-2-amino-3-(benzhydrylthio)-N-((2S,3S,4R,5R)-2-(chloromethyl)-5-(6-(dimethylamino)-9H-purin- 9-yl)-4-hydroxytetrahydrofuran-3-yl)propenamide (**25)**

The crude product obtained by following general procedure B as described above, was purified by flash chromatography utilizing Teledyne ISCO’s Combiflash^®^ R_f_ system equipped with normal phase RediSep (silica) columns with methanol/chloroform eluent mixture (neat CHCl_3_ for 8 min, to 20% MeOH/CHCl_3_ gradual gradient over the next 31 min) furnished compound **25** as xx solid in 7.3 mg (27%) yield; R_f_ 0.35 (MeOH:CHCl_3_/1:19); **^1^H-NMR** (600 MHz, CD_3_OD): 8.136 (2H, d, J = 1.09 Hz), 7.40-7.28 (m, 4H), 7.23-7.15 (m, 4H), 7.14-7.08 (m, 2H), 5.971 (1H, d, J = 2.11 Hz), 5.203 (s, 1H), 4.653 (1H, dd, J = 8.45, 5.98 Hz), 4.503 (1H, dd, J = 5.97, 2.07 Hz), 4.226 (1H, ddd, 8.34, 5.18, 3.01), 3.882 (1H, dd, J = 12.25, 2.83 Hz), 3.750 (1H, dt, J = 7.92, 4.37 Hz), 3.45-3.40 (6H, m), 2.673 (1H, dd, J = 13.35, 5.60 Hz), 2.579 (1H, dd, 13.35, 6.74 Hz); **^13^C-NMR** (101 MHz, CD_3_OD) 174.34, 154.74, 151.89, 149.51, 141.78, 141.01, 137.10, 128.60, 128.21, 128.18, 126.68, 119.90, 90.27, 81.00, 73.81, 58.52, 57.63, 51.66, 43.79, 37.61, 31.60; ESI-MS *m/z* [M + H]^+^ 582.2.

### Biology

#### Cloning and purification of Puromycin−Sensitive Aminopeptidase (PSA)

PSA was amplified from human cDNA (gene name NPEPPS), cloned using the InFusion enzyme system and purified as described before [26]. Briefly, the expression vector, pEXP−HisMBP−NPEPPS expresses TEV cleavable PSA with an N−terminal His−tagged MBP solubility enhancer. For PSA expression, pEXP−HisMBP−NPEPPS was transformed into BL21 STAR (DE3) *E. coli* cells. Cultures were induced with 0.5 mM IPTG at an OD_600_ of 0.6 with shaking overnight at 16 °C. Cell pellets were resuspended in lysis buffer (50 mM HEPES, 300 mM NaCl, 10 mM imidazole, pH 8.0) and sonicated. The lysate was centrifuged at 50,000×g for 10 min and to the supernatant 1 mL of 50% Ni−NTA (Qiagen) was added and incubated for 1 h at 4 °C. The Ni-NTA was washed with 16 mL of wash buffer (50 mM HEPES, 300 mM NaCl, 20 mM imidazole, pH 8.0) and eluted with 2.5 mL of elution buffer (50 mM HEPES, 300 mM NaCl, 250 mM imidazole, pH 8.0) with the flow through collected and desalted on a PD−10 column into TEV cleavage buffer (50 mM Tris·HCl, pH 8.0, 0.5 mM EDTA, 300 mM NaCl, 1 mM DTT). The His-MBP tag was cleaved with His-tagged TEV protease, and both were removed with Ni-NTA resulting in cleaved PSA protein. The cleaved protein was concentrated with a 3000 Da molecular weight cutoff spin filter.

#### Dose–Response Studies of PSA

PSA inhibition assays were performed as described previously [26]. Briefly, 40 nM PSA, 0.8 µL of inhibitor and 25 mM Tris, pH 7.5 were incubated on a black 384–well plate at a final volume of 80 µL. The addition of alanine-4-methoxy-2-naphthylamide (Ala-4-MNA) substrate (Sigma) at a final concentration of 190 µM starts the reaction. Peptide cleavage and resulting accumulation of 4–MNA was measured using fluorescence, exciting at 340 nm and reading at 425 nm at room temperature for 30 min on a Molecular Devices SpectraMax i3 plate reader. IC_50_ values were determined by fitting steady-state velocities vs. inhibitor concentration to the sigmoidal concentration–response curve (variable slope) using GraphPad Prism.

#### Does-Response Studies of APN

APN assays were performed as described previously [26]. APN was purchased from R&D Systems. Amino-4-methylcoumarin (Ala–AMC) was purchased from Bachem. Briefly, 0.1 µg/µL APN, 1 µL inhibitor and 50 mM Tris buffer, pH 7.5 were incubated in a black 384–well plate at a final volume of 100 µL. The addition of Ala–AMC substrate at a final concentration of 100 µM starts the reaction. Peptide cleavage and resulting accumulation AMC was measured using fluorescence, exciting at 340 nm and reading at 425 nm at room temperature for 30 min on a Molecular Devices SpectraMax i3 plate reader. IC_50_ values were determined by fitting steady state velocities vs. inhibitor concentration to the sigmoidal concentration–response curve (variable slope) using GraphPad Prism.

#### Tail-Flick Assay

The analgesic potency of the lead PSA inhibitor was tested in tail-flick assay using Columbus Instruments tail-flick analgesiometer. Compound **19** was dissolved in saline using 2 equivalents of aqueous HCl (forming hydrochloride salt). Eight-week-old male CF-1 mice were used for these experiments. Mice were gently restrained in a cloth before being exposed to the tail-flick beam on the dorsal surface of the tail (about 15 mm from the tip of the tail). The time required for withdrawal of the tail was recorded as latency. The intensity of the heat source was adjusted to achieve baseline measurement of 2−4 s. The compound was administered by i.c.v. route (50 or 100 µg/mouse) alone or in combination with Met-Enk (50 µg/mouse in saline). For experiments with naloxone, subcutaneous naloxone (5 mg/kg in water) was injected 5 minutes after the i.c.v. injection. The tail-flick response was measured at 15, 30, 45, and 60 min after drug administration. Maximum latency was recorded as 10 s to prevent tissue injury.

#### Hot-Plate Method

Eight-week-old male CF1 mice were administered with compounds as described above in tail-flick assay. Temperature of the hot plate which is a cylindrical glass bath was maintained thermostatically at 55 °C by circulating hot water. Response time of mice for hind-paw shake or lift or hind-paw lick or jump from the platform was recorded as latency. Baseline latencies was recorded at mean reaction time of 12 s and mice with higher reaction times were excluded as non-responders. The reaction times of mice injected with test compounds were recorded at 15 min post administration. A cutoff time of 60 s was used as the maximum latency to avoid tissue injury.

Analgesic response by tail-flick and hot plate assay are represented as the percentage maximum possible effect (%MPE) calculated using %MPE = (test − baseline)/(cutoff −baseline) × 100, where test is the observed latency after compound treatment, baseline is the latency prior to the treatment, and cutoff is the time used as the end of the test i.e. 10 s for tail flick and 60 s for hot plate assay. Statistical analysis was conducted by One-way ANOVA using Tukey’s multiple comparison test.

#### Mass Spectrometry-based analysis of enkephalin levels

Coronal slices from wild type mice (300 µm thick) submerged in 100 µL artificial cerebrospinal fluid (aCSF) ± peptidase inhibitors were stimulated with KCl (50 mM) for 20 min in the presence and absence of PSA inhibitor (final concentration 10 µM). The extracellular fluid was collected for analysis of opioid peptides levels using LC-MS/MS protocol described previously [43]. Briefly, samples after desalting through C-18 stage tips (SP301, Thermo Scientific), and concentration under vacuo were reconstituted in water:acetonitrile:formic acid (98:2:0.1). Samples (5 µL) were injected onto an analytical C18 reverse phase column (Phenomenex, Torrance, CA) using gradient elution with 0.1% FA in water and 0.1% FA in ACN as mobile phase at 1 µL/min flow rate. Previously established mass transitions for neuropeptides were utilized for detection on a TSQ Quantiva Triple Quadrupole (Thermo Scientific) in positive nanospray ionization mode [43]. Accuracy of peptide detection was verified by manual inspection and integration of target peaks. Statistical analysis was conducted using non-parametric Mann-Whitney one-tailed U-test.

#### Plasma stability assessment

The test compound (1 µM in incubation) was spiked into pooled human or mouse plasma and incubated at 37°C. The incubations were performed in triplicate. At 0, 1, 3, 6 and 24 h, 40 µL of incubation mixture were removed and immediately quenched by 120 µL of acetonitrile containing internal standard (0.05 µM). The quenched samples were centrifuged at 14,000 rpm for 10 min at 4°C. The supernatants were stored at -20°C until submitted for analysis by LC-MS/MS. Plasma stability of the test compounds were evaluated by monitoring the disappearance of test compound over a 24 h time period. The peak area ratios of analyte versus internal standard were used to calculate the remaining percentage at each time point using Microsoft excel.

#### Microsomal stability assay

The test compound (5 µM in incubation) was pre-incubated in 0.1 M potassium phosphate buffer (pH 7.4) containing human or mouse liver microsomes (0.5 mg/mL in incubation) at 37°C for 5 min. The reaction was initiated by the addition of NADPH (1 mM in incubation) and incubated at 37°C in water bath. The incubations were performed in triplicate. A negative control was performed in parallel in the absence of NADPH to reveal any chemical instability or non-NADPH dependent enzymatic degradation for each test compound. A positive control, verapamil (1 µM in incubation), was also performed to confirm the proper functionality of the incubation system. At 0, 5, 15 and 30 min, 50 µL of the incubation mixture was removed and immediately quenched by 150 µL of acetonitrile containing an appropriate amount of internal standard (0.1 µM for **19**, 0.5 µM for verapamil). The quenched samples were sat on ice for 5 min and centrifuged at 14,000 rpm for 10 min at 4°C. The supernatants were stored at -20°C until submitted for analysis by LC-MS/MS. Microsomal stability of the test compounds was evaluated by monitoring the disappearance of the test compound over a 30 min time period. The peak area ratios of analyte versus internal standard were used to calculate the remaining percentage at each time point using Microsoft Excel. The natural logarithm of the remaining percentage is plotted against time and the gradient of the line is used to determine turnover rate constant k (min^-1^), t1/2 and intrinsic clearance (CL_int_).

k (min^-1^)= (-gradient)

t1/2 (min) = 0.693/k

V (µL/mg) = volume of incubation/protein in incubation

CLint (µL/min/mg protein) = V*k

The samples were injected in an Agilent 1260 HPLC coupled to an AB Sciex QTRAP 5500 mass spectrometer and separated using a Phenomenex Kinetex C18 column (50 mm × 2.1 mm, 2.6 µm) with a mobile phase of 0.1% formic acid in water with 2% acetonitrile (mobile phase A) and 0.1% formic acid in acetonitrile (mobile phase B) at a flow rate of 0.5 ml/min. The analytes were eluted with a gradient as follows: from 0 to 0.5 min, 5%B (v/v); from 0.5 to 1 min, 5–95% B (v/v); from 1 to 3 min, 95% B (v/v); from 3 to 3.2 min, 95–5% B (v/v); and from 3.2 to 6 min, 5% B (v/v). All samples were analyzed by an electrospray ionization source in positive mode. The optimized source and gas parameters were as follows: curtain gas, 30 psi; CAD gas, medium; ion spray voltage, 5000 V; temperature, 700 °C; gas 1, 60 psi; gas 2, 50 psi. Multiple reaction monitoring (MRM 1) was conducted by monitoring the following transition: m/z 582.2→419.2 (compound **19**); m/z 455.3→165.1 (verapamil); m/z 569.3→434.3 (internal standard).

MRM transitions of potential metabolites were predicted using LightSight^®^ software. An information dependent acquisition (IDA) method was created using the predicted MRM transitions as survey scan and the enhanced product ion (EPI) scan as dependent scans.

IDA criteria: The two most intense peaks which exceeds 1000 cps after dynamic background subtraction of survey scan will be selected and subjected to EPI scans.

Metabolites Profiling: Two samples, which represented the 0 and 5 min time points in microsomal stability assay, were injected into LC-MS/MS with a volume of 5 µL and analyzed by the IDA method described above. The metabolites profiling was performed in both species, mouse and human liver microsomes. Data obtained in the metabolites profiling was analyzed using the LightSight® software.

#### Docking of PSA targeting compounds

Molecular modeling was performed using the Schrödinger Maestro 2022-2 [26, 27]. The crystal structure of Puromycin-sensitive aminopeptidase (PDB code 8SW0) was used as a starting point [23]. This model was subjected to Protein Preparation Wizard for the addition of missing protein hydrogen atoms and removal of water molecules beyond 5 Å. This was followed by the addition of missing side chains and loops using Prime and optimization of the hydrogen bonding network using PROPKA (pH 7.0). The protein structure was then minimized using OPLS4 force field [48] and receptor grid was generated. The active site in the receptor grid covered all amino acid residues within a 25 Å^3^ box with coordinates X: 29.974, Y: 83.117 and Z: 2.974. The flexibility of the grid was ensured by incorporating van der Waals radii of nonpolar atoms in the receptor and partial charge cutoff of 0.25. Puromycin, **19** and **22**, along with the corresponding 5′-hydroxy (OH) analogs (compounds **20** and **23**, respectively) were sketched using Maestro 2D-sketcher and subjected to LigPrep to generate conformers incorporating metal states in the pH range of 7 ± 2 to be utilized for the docking process.

Ligand docking was performed in Extra Precision (XP) mode using Glide [27]. All the ligands were incorporated as flexible entities, and strain correction terms as well as post-docking minimization were performed on the generated poses.

## ASSOCIATED CONTENT

### Supporting Information

Additional data on docking studies and compound characterization including ^1^H and ^13^C NMR of final compounds and HPLC data of the compound tested in animals. This material is available free of charge via the Internet at XXXX.

## AUTHOR INFORMATION

### Author Contributions

All authors have given approval to the final version of the manuscript.

### Institutional Review Board Statement

All experimental protocols involving mice were in strict adherence to the NIH Animal Care and Guidelines and were approved by the Institutional Animal Care and Use Committee (IACUC) at the University of Minnesota, Minneapolis, MN (protocol code 2204-39921A, last approved on 8 December 2022; Expiration Date 28 June 2025).

## ACKNOWLEDGMENT

The authors would like to thank Dr. Harrison VanKoten and Ms. Christine Dreis for technical help with characterization of the compounds.

## ABBREVIATIONS

PSA: puromycin sensitive aminopeptidase
APN: aminopeptidase N
NEP: neprilysin
ACE: angiotensin converting enzyme
PAN: puromycin aminonucleoside
OUD: opioid use disorder
Met-Enk: Metenkephalin
Leu-Enk: Leu-Enkephalin
MERF: Met-Enkephalin-Arg-Phe
MERGL: Met-enkephalin-ArgGly-Leu
DENKIs: dual enkephalinase inhibitors
TFA: trifluoroacetic acid
FA: formic acid
SAR: structure activity relationship
MPE: maximum possible effect
TLC: thin layer chromatography
HPLC: high performance liquid chromatography
LC-MS/MS: liquid chromatography tandem mass spectrometry.

