## Supporting Information for "Modulation of endogenous opioid signaling by inhibitors of puromycin sensitive aminopeptidase"

### Table of Contents

**Figure S1. Ligand interaction diagram of compound 20** **S2**
**Figure S2. Ligand interaction diagram of compound 23** **S2**
**Table S1. Binding energy scores of PSA inhibitors** **S3**
**$^{1}$H NMR and $^{13}$C NMR spectra of compound 18** **S4**
**$^{1}$H NMR and $^{13}$C NMR spectra of compound 19** **S5**
**HPLC chromatogram of compound 19** **S6**
**$^{1}$H NMR and $^{13}$C NMR spectra of compound 21** **S7**
**$^{1}$H NMR and $^{13}$C NMR spectra of compound 22** **S8**
**$^{1}$H NMR and $^{13}$C NMR spectra of compound 24** **S9**
**$^{1}$H NMR and $^{13}$C NMR spectra of compound 25** **S10**

**Figure S1.** Molecular docking analysis of compound **20**. Left image shows fitting of the inhibitor within the binding site, while right image shows 2D interactions of the inhibitor with the PSA active site Zn(II) and surrounding residues. Orientation of compound **20** is similar to that of compound **19**, which was directed by purine ring binding to Zn(II) and accommodation of the aromatic amino acid residue in the hydrophobic pocket.

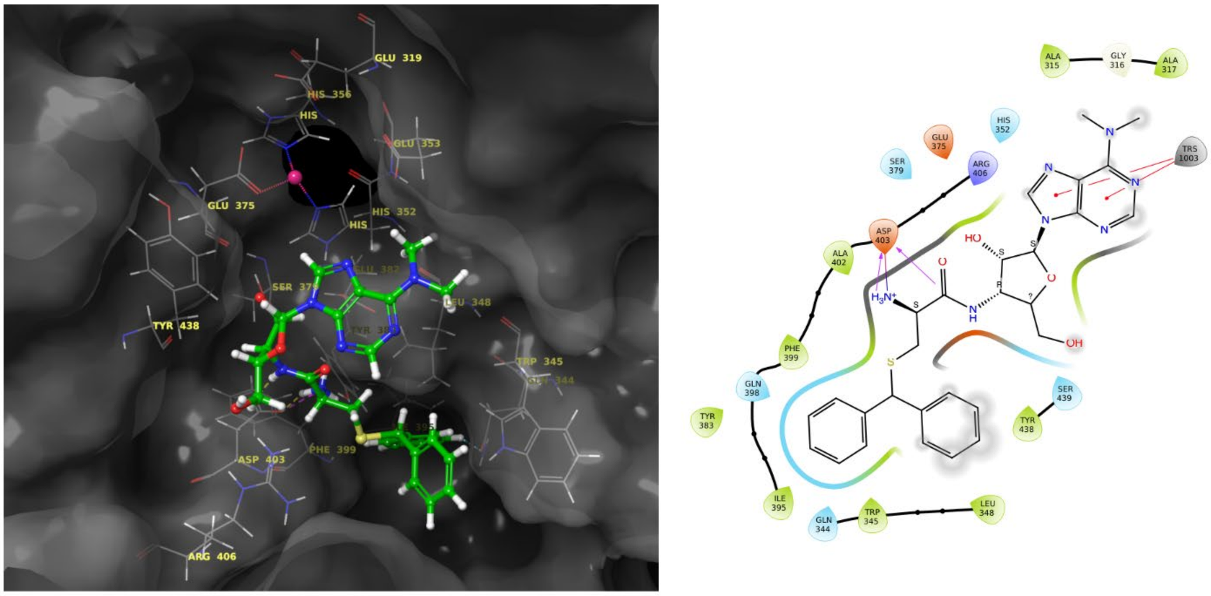


**Figure S2.** Molecular docking analysis of compound **23**. Two different conformations were obtained after docking of compound **23**. Shown in (A) is the overlap of the two conformations. (B) Binding of the purine ring in the hydrophobic pocket, away from the active site Zn(II), improved binding energy. (C) Purine ring in the proximity of Zn(II) resulted in binding conformation similar to the corresponding chloro compound **22**, and offered comparable active site interactions and docking score.

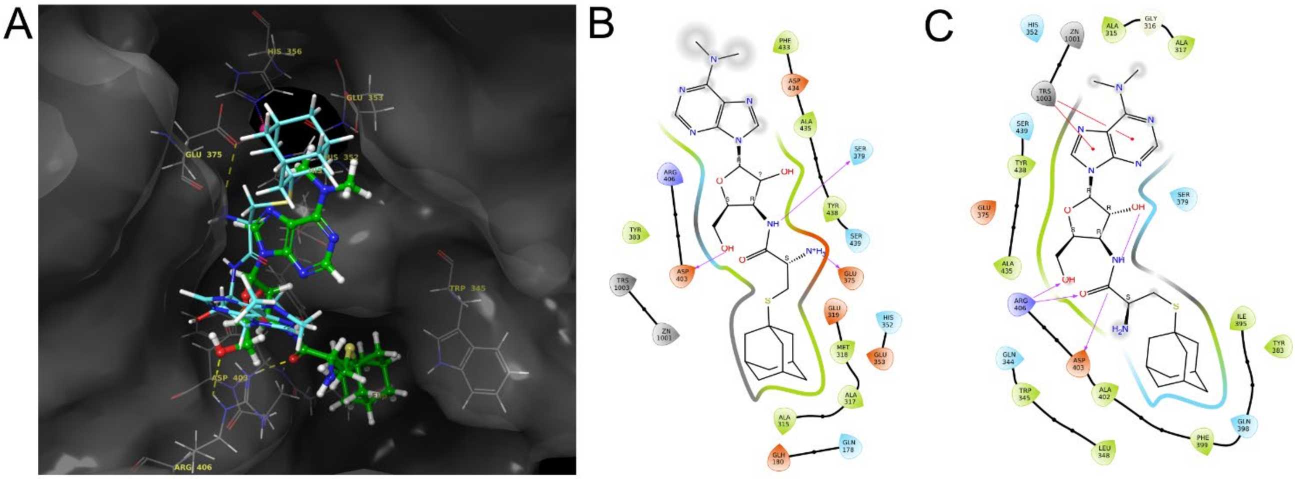

**Table S1: Binding energy scores of the compounds with and without Zn(II) as constraint**

| Name of Compound | Zn ion as constraint (Y/N) | 5′-Chloride group present (Y/N) | Binding energy score (kcal/mol) |
|---|---|---|---|
| Puromycin | N | N | -8.390 |
| 19 | N | N | -7.578 |
| 22 | N | N | -6.060 |
| 20 | N | Y | -8.050 |
| 23 | N | Y | -7.212 |
| Puromycin | Y | N | -8.548 |
| 19 | Y | N | -7.627 |
| 22 | Y | N | -7.457 |
| 20 | Y | Y | -7.984 |
| 23 | Y | Y | -7.946 |

¹H NMR (400 MHz, $CD_3OD$) and 13C NMR (101 MHz, $CD_3OD$) spectra of **18**

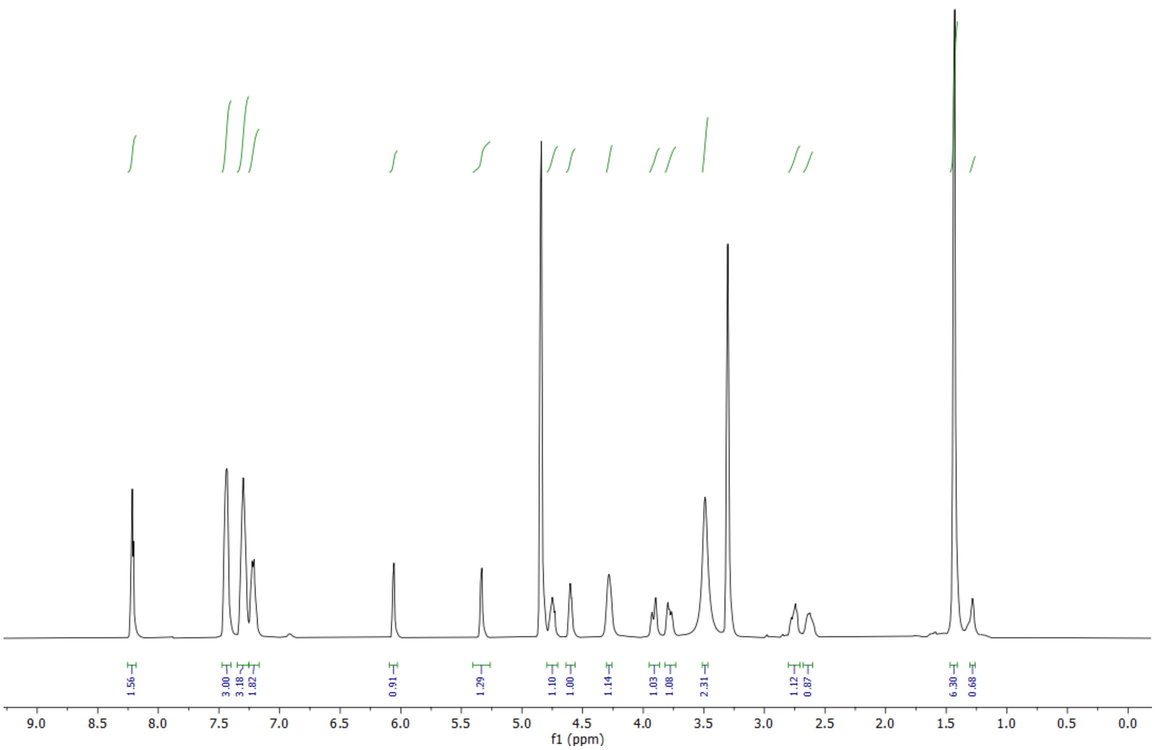


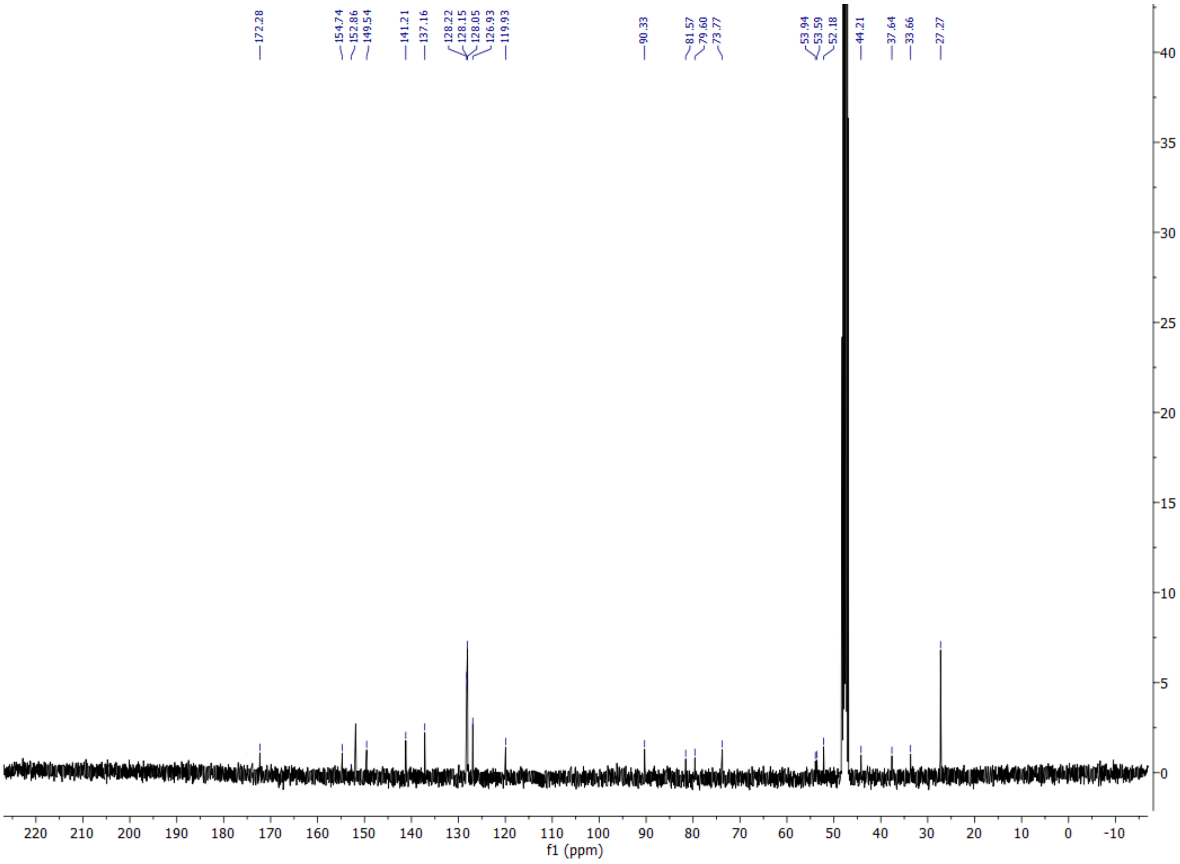

$^{1}H$ NMR (400 MHz, $CD_3OD$) and 13C NMR (101 MHz, $CD_3OD$) spectra of **19**

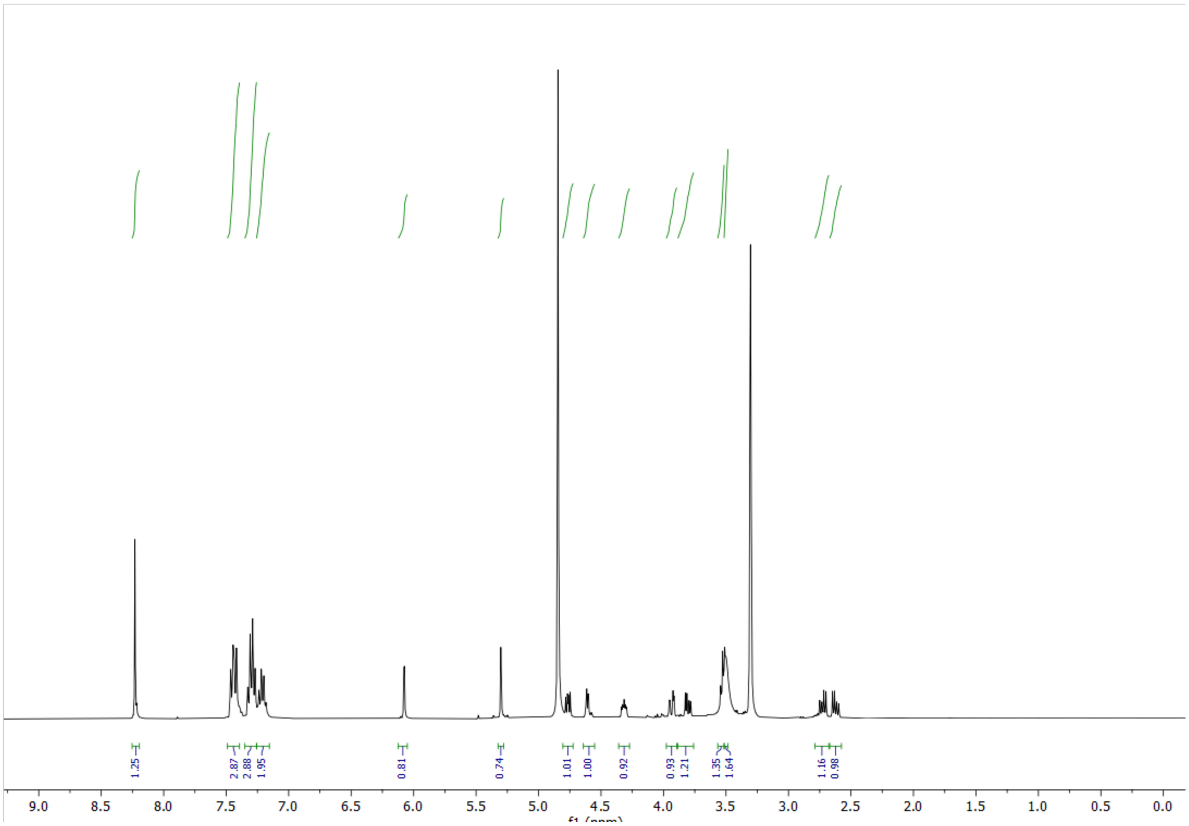

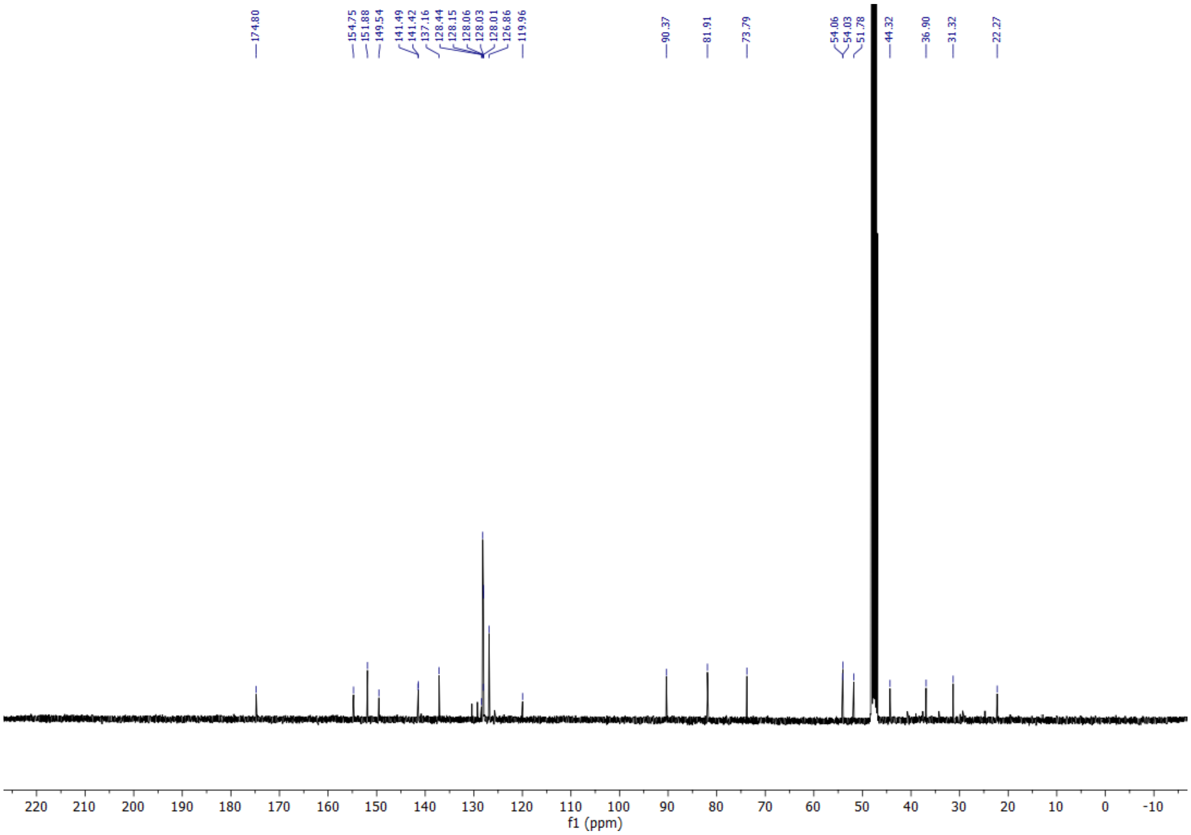

HPLC Chromatogram of compound **19**

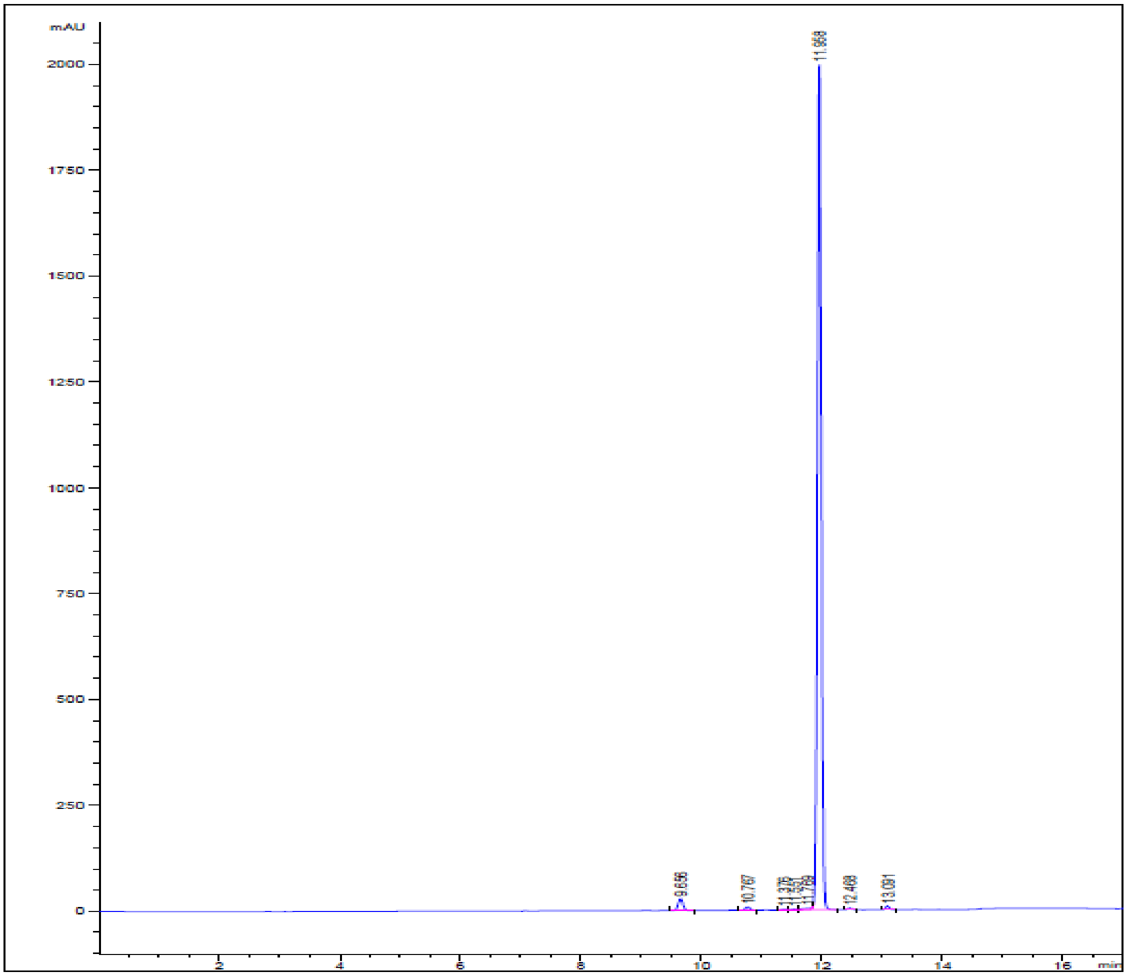

| Peak # | RetTime (min) | Width (min) | Area (mAU*s) | Height (mAU) | Area % |
|---|---|---|---|---|---|
| 1 | 9.656 | 0.08 | 151.16 | 28.01 | 1.59 |
| 2 | 10.767 | 0.08 | 40.11 | 7.79 | 0.42 |
| 3 | 11.376 | 0.09 | 10.94 | 1.79 | 0.12 |
| 4 | 11.551 | 0.09 | 12.77 | 1.86 | 0.14 |
| 5 | 11.769 | 0.14 | 38.05 | 3.59 | 0.40 |
| 6 | 11.958 | 0.07 | 9154.05 | 1996.81 | 96.77 |
| 7 | 12.468 | 0.07 | 15.05 | 3.35 | 0.16 |
| 8 | 13.091 | 0.07 | 37.11 | 8.50 | 0.39 |

$^{1}H$ NMR (400 MHz, $CD_3OD$) and 13C NMR (101 MHz, $CD_3OD$) spectra of **21**

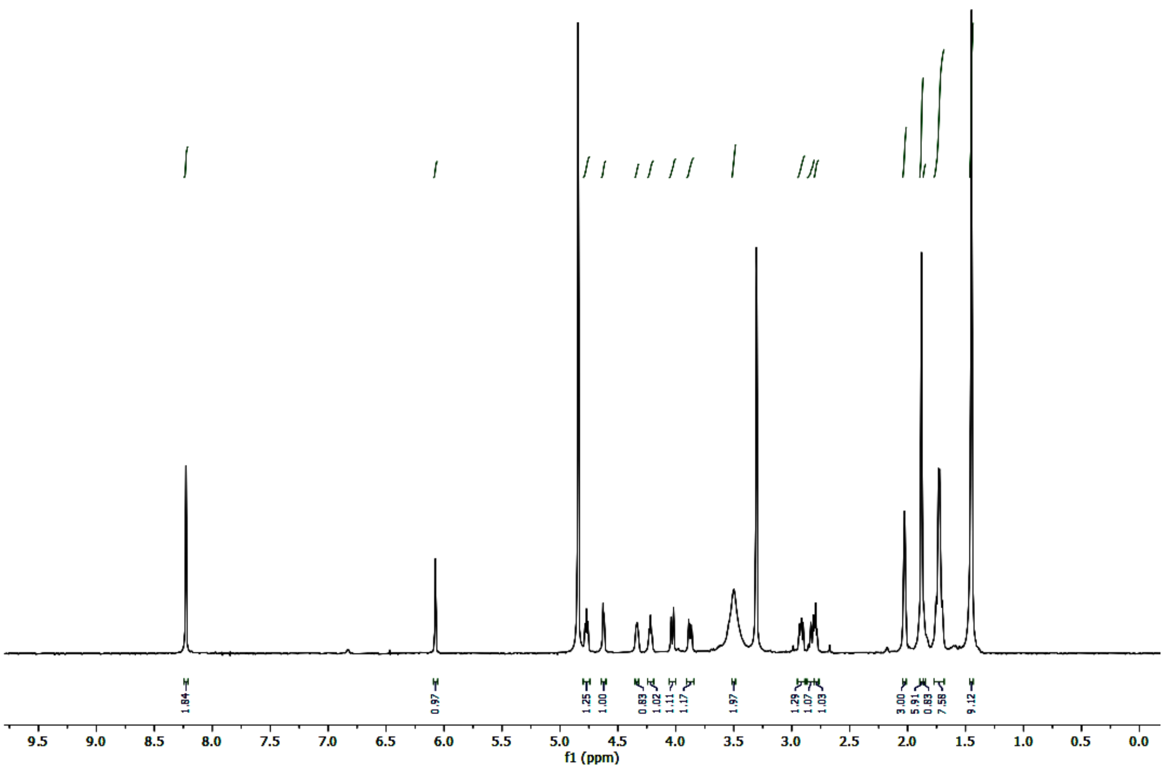


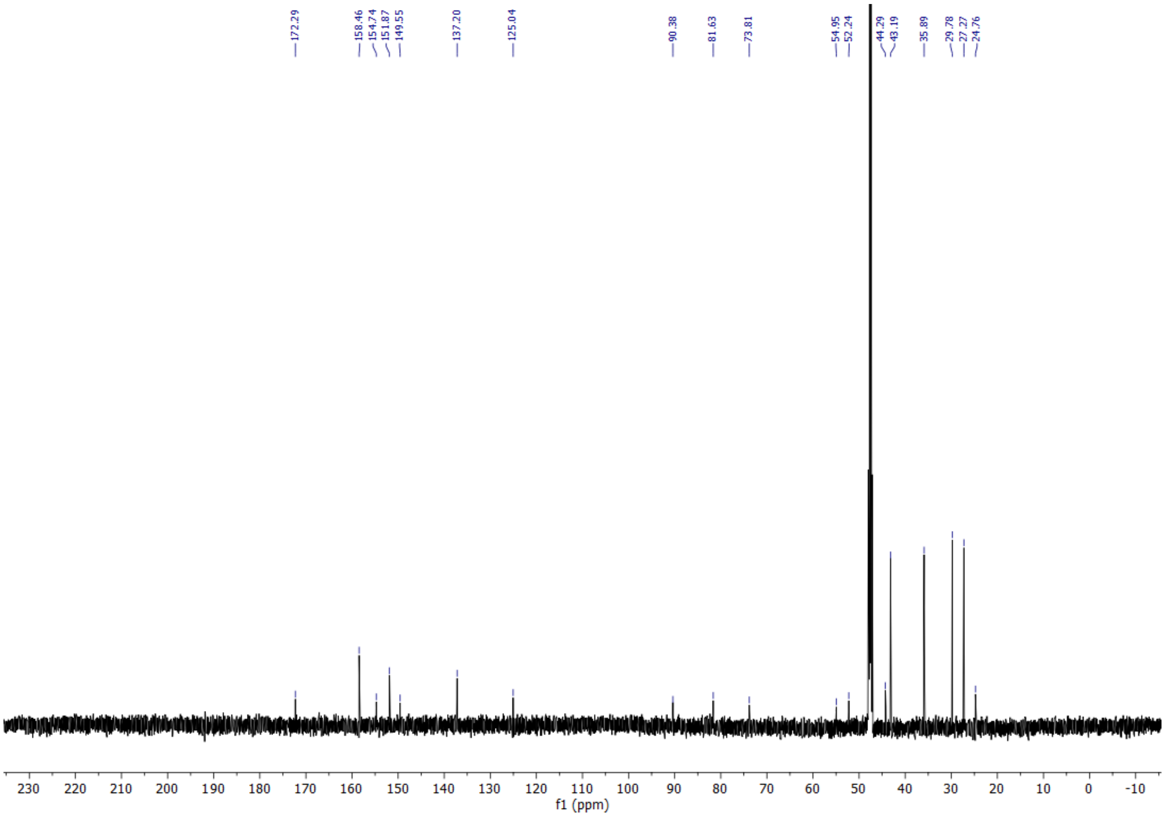

[1]H NMR (400 MHz, $CD_3OD$) and 13C NMR (101 MHz, $CD_3OD$) spectra of **22**

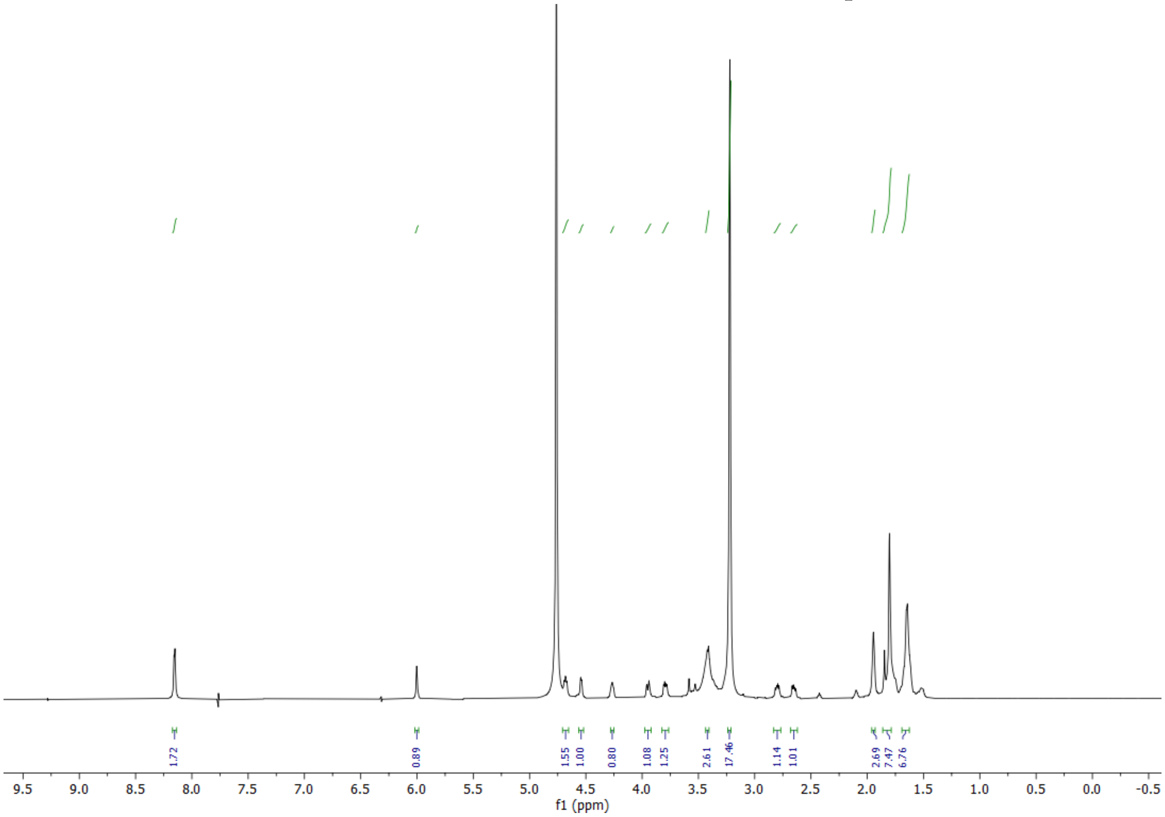


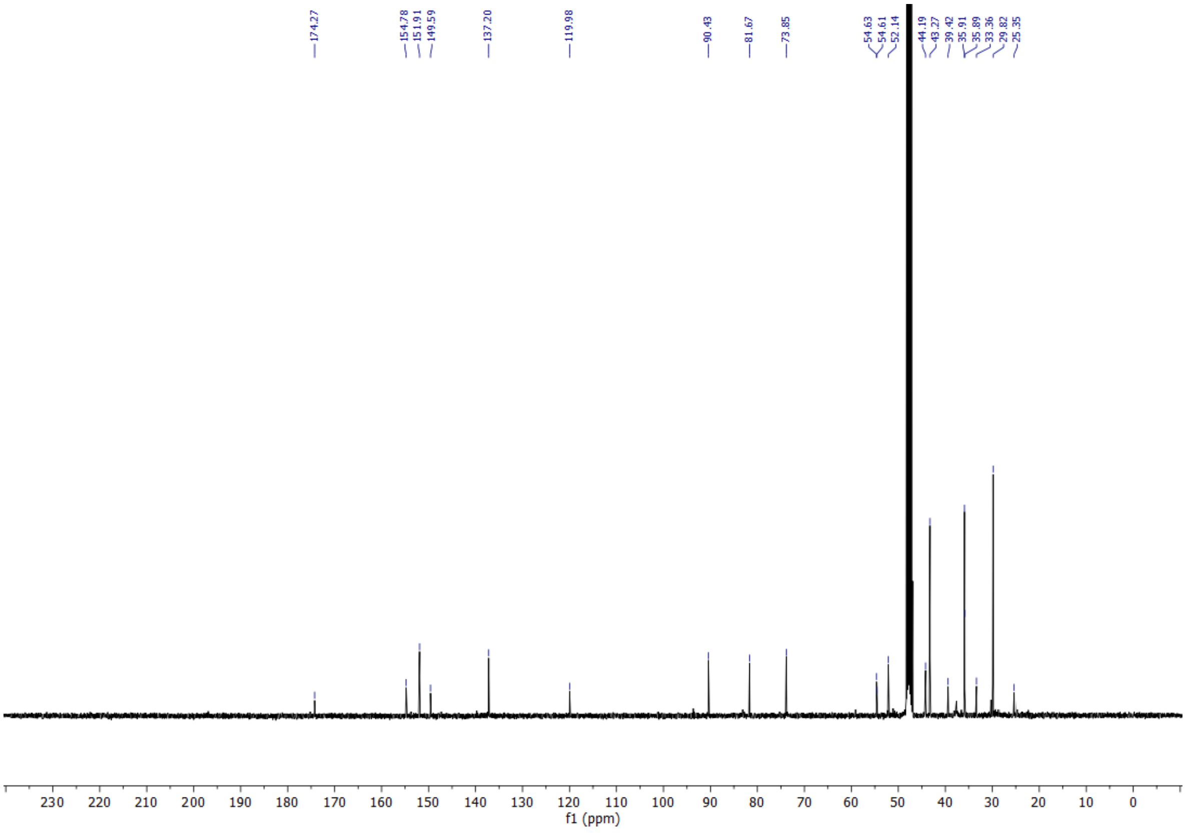

[1]H NMR (400 MHz, $CD_3OD$) and 13C NMR (101 MHz, $CD_3OD$) spectra of **24**

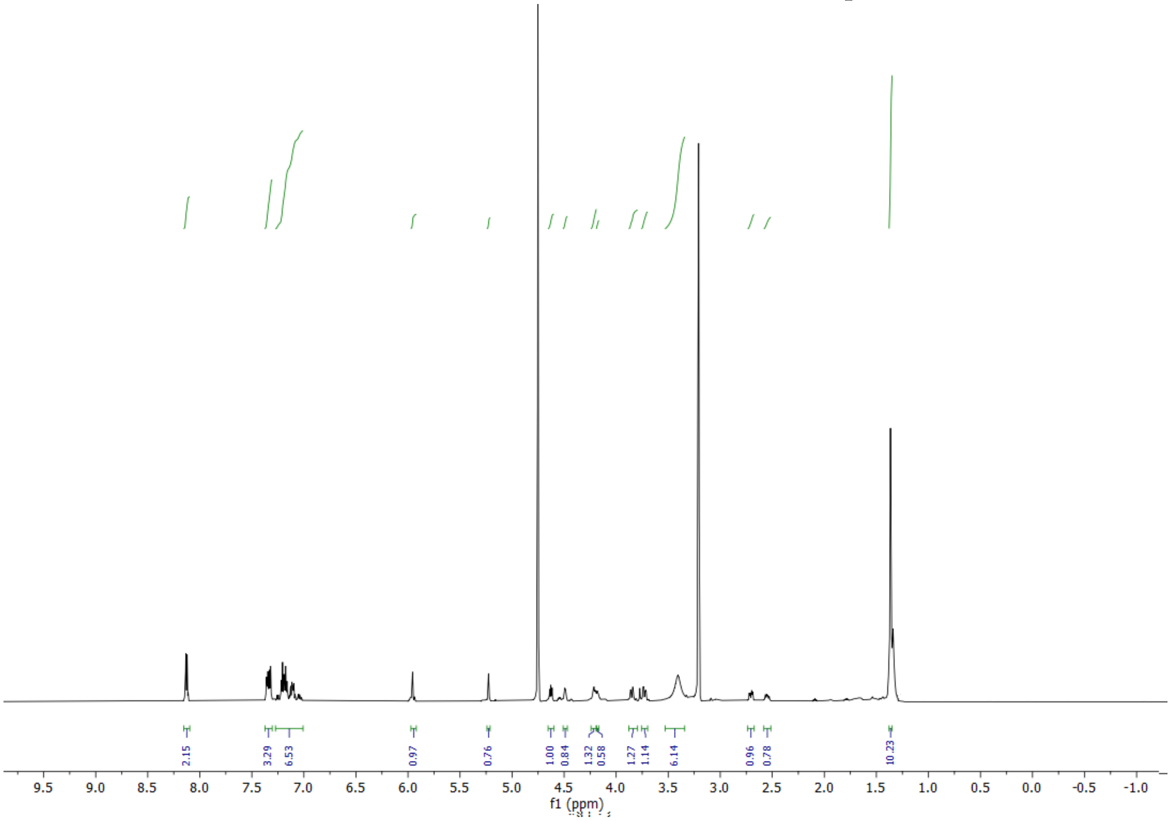

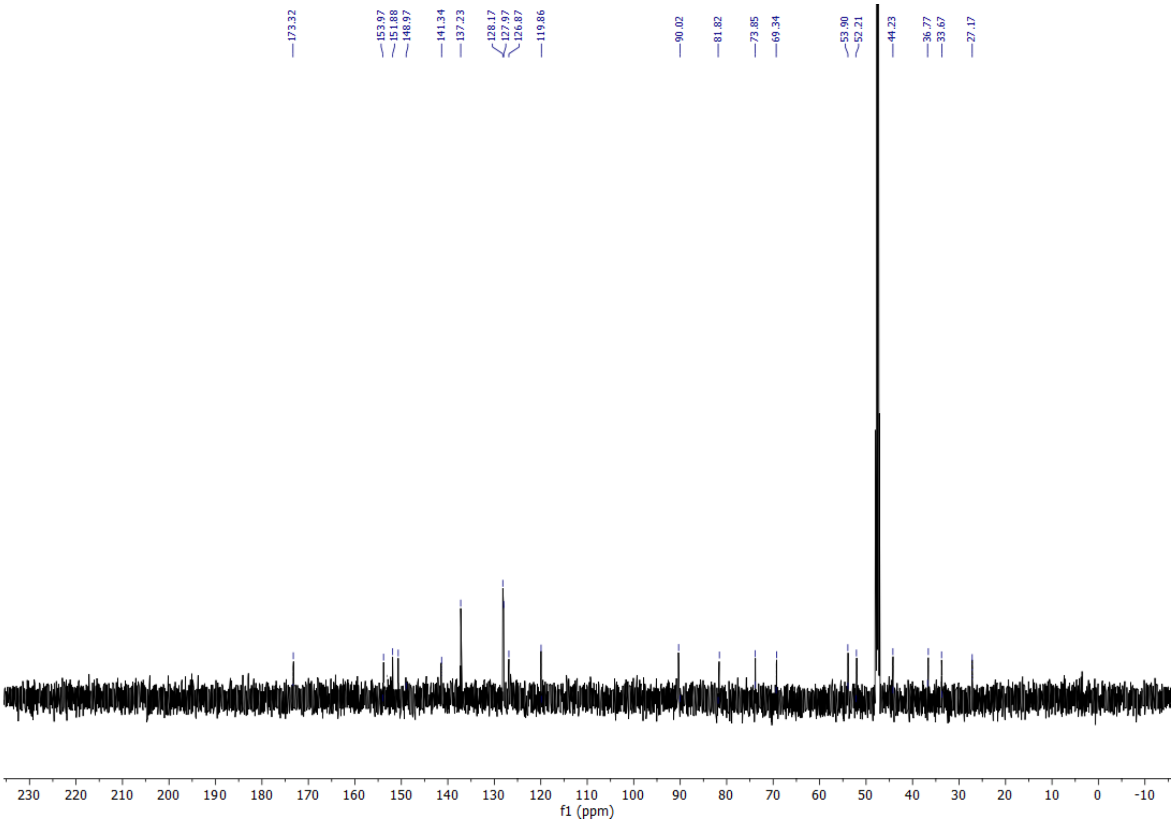

[1]H NMR (400 MHz, CD3OD) and 13C NMR (101 MHz, CD3OD) spectra of **25**

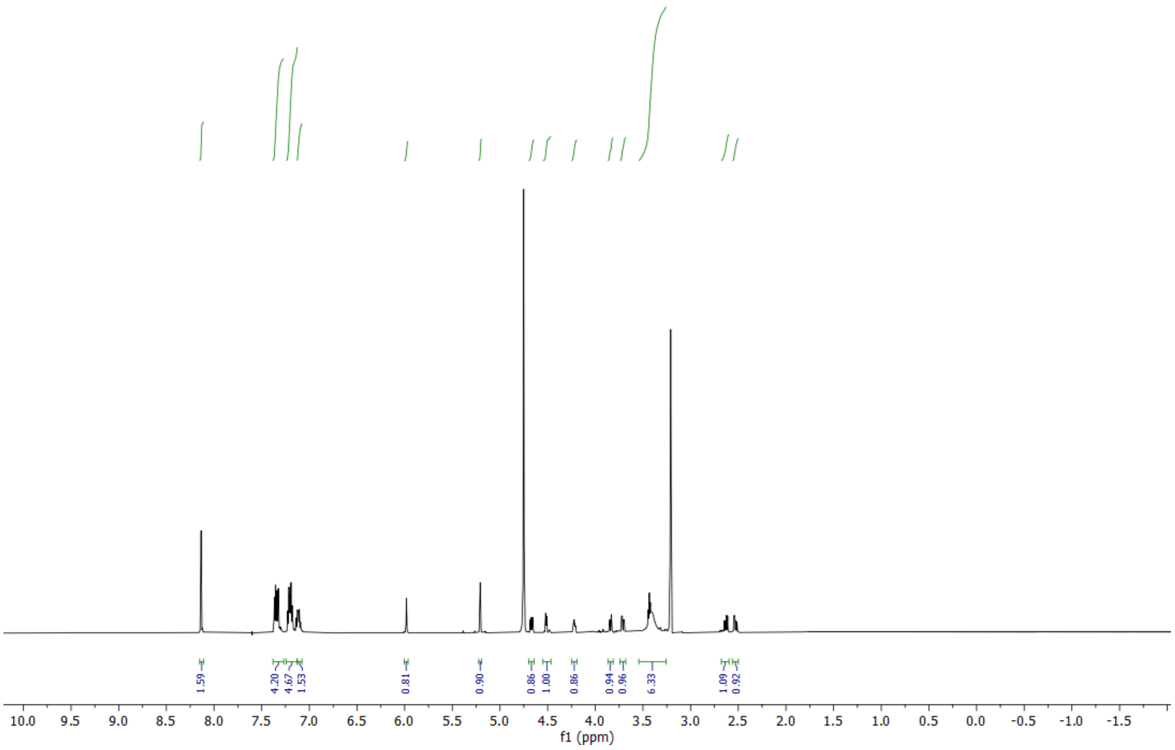

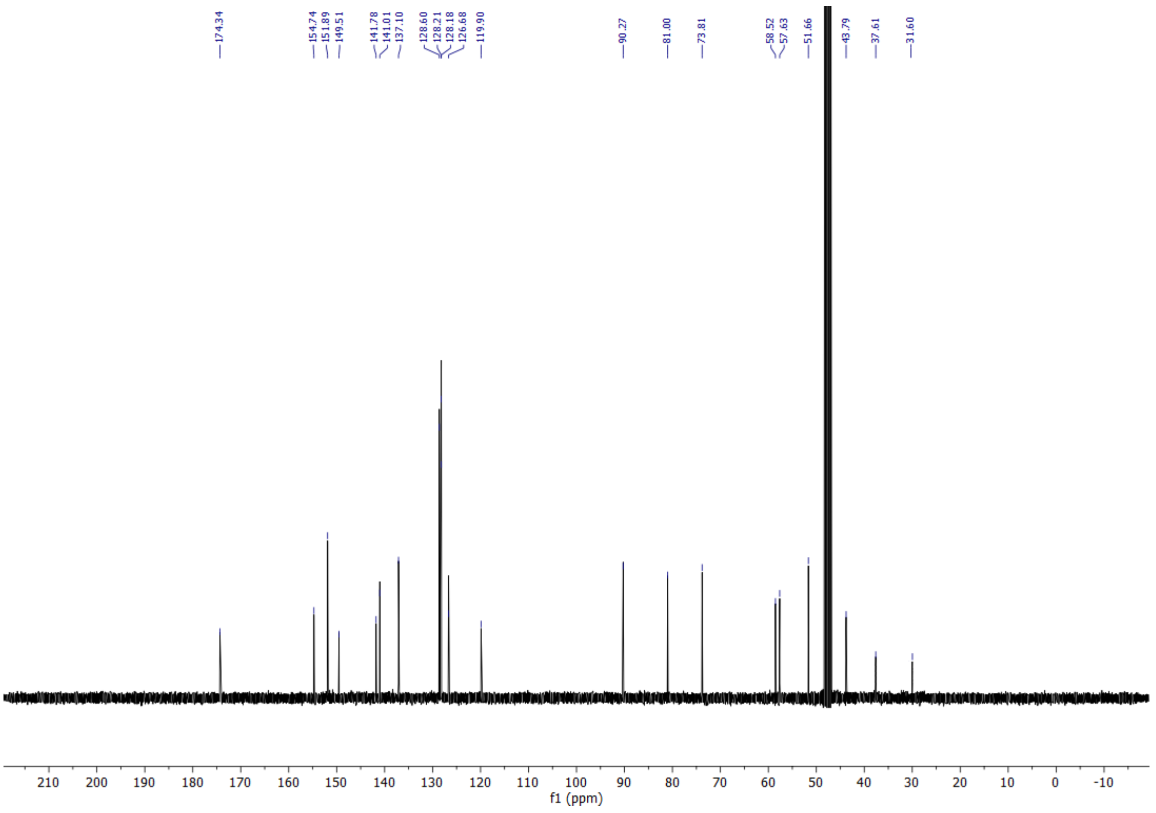
